# MITF Targets in Gastrointestinal Stromal Tumors: Implications in Autophagy and Extracellular Vesicle Secretion

**DOI:** 10.1101/2024.09.10.612253

**Authors:** Elizabeth Proaño-Pérez, Eva Serrano-Candelas, Mario Guerrero, David Gómez-Peregrina, Carlos Llorens, Beatriz Soriano, Ana Gámez-Valero, Marina Herrero-Lorenzo, Eulalia Martí, César Serrano, Margarita Martin

**Affiliations:** Biochemistry and Molecular Biology Unit, Biomedicine Department, Faculty of Medicine and Health Sciences, University of Barcelona, Barcelona, Spain; Clinical and Experimental Respiratory Immunoallergy (IRCE), Institut d’Investigacions Biomediques August Pi i Sunyer (IDIBAPS), Barcelona, Spain; Faculty of Health Sciences, Technical University of Ambato, Ambato-Ecuador; ProtoQSAR SL, Centro Europeo de Empresas Innovadoras (CEEI), Parque Tecnológico de Valencia, 46980 Valencia, Spain; Biotechvana, Parc Científic Universitat de València, Valencia, Spain; Sarcoma Translational Research Program, Vall d’Hebron Institute of Oncology (VHIO), Hospital Universitario Vall d’Hebron, Vall d’Hebron Barcelona Hospital Campus, C/ Natzaret, 115-117, 08035, Barcelona, Spain; Department of Medical Oncology, Vall d’Hebron University Hospital, Barcelona, Spain

**Author notes:** **CORRESPONDENCE:** M. Martin, Biochemistry Unit, Biomedicine Department, Faculty of Medicine, University of Barcelona, Carrer de Casanova 143, E-08036 Barcelona, Spain. **Acronyms** Chromatin Immunoprecipitation sequencing (ChIP-seq); Enzyme commission number (EC); False Discovery Rate (FDR); Fraction of reads in peaks (FRiP); Gastrointestinal Stromal Tumors (GIST); Gene ontologies (GO); Markov Cluster (MLC); Microphthalmia-associated Transcription Factor (MITF); Protein-protein interaction network (PPI Network); Quantitative PCR (qPCR).

**Keywords:** MITF / ChIP-seq / Autophagy / Extracellular vesicles/ Gastrointestinal stromal tumors (GIST)

## Abstract

Previous studies have identified Microphthalmia-associated Transcription Factor (MITF) involvement in regulating Gastrointestinal Stromal Tumors (GIST) growth and cell cycle progression. This study uses Chromatin Immunoprecipitation combined with high-throughput sequencing (ChIP-seq) and RNA sequencing to explore MITF-modulated genes in GIST. Our findings reveal that MITF regulates genes involved in lysosome biogenesis, vesicle generation, autophagy, and mTOR signaling pathways. Comparative transcriptome analysis following MITF silencing in GIST cells shows differential enrichment in mTOR signaling, impacting tumor growth and autophagy. In the context of cancer, the interplay between autophagy and extracellular vesicle release can influence tumor progression and metastasis. We examined MITF’s role in autophagy and extracellular vesicle (EV) production in GIST, finding that MITF overexpression increases autophagy, as shown by elevated LC3II levels while silencing MITF disrupts autophagosome and autolysosome formation. Despite no significant changes in EV size or number, MITF silencing notably reduces KIT expression in EV content. KIT secretion in EVs has been linked to GIST metastasis, suggesting that MITF is a crucial target for managing tumor growth and metastasis in GIST.

## INTRODUCTION

The microphthalmia-associated transcription factor (MITF) belongs to the MiT/TFE family. In humans, it comprises four evolutionarily conserved members: microphthalmia-associated transcription factor (MITF), transcription factor EB (TFEB), TFE3, and TFEC (Steingrímsson *et al*, 2004). They can form homodimers and heterodimers and bind, through the basic domain, to the regulatory regions of their target genes that contain the 6-base pair CANNTG motif (E-box) (Hemesath *et al*, 1994; Pogenberg *et al*, 2020). MiT/TFE proteins are regulated at transcriptional and posttranscriptional levels (Puertollano *et al*, 2018; Ngeow *et al*, 2018; Goding & Arnheiter, 2019) and are involved in an exhaustive list of physiological functions, including cell survival, differentiation, senescence, metabolism, response to endoplasmic reticulum (ER) stress, mitochondrial and DNA damage, oxidative stress, innate immunity and inflammation, longevity, neurodegeneration, and cancer (Goding & Arnheiter, 2019; La Spina *et al*, 2021; Raben & Puertollano, 2016; Cortes & La Spada, 2019; Bahrami *et al*, 2020; Irazoqui, 2020; Wang *et al*, 2020).

Microphthalmia transcription factor (MITF), which plays a significant role in oncogenesis, has been widely investigated in melanoma. Indeed, MITF is a master regulator of multiple aspects of melanocyte function. MITF-M is the specific and highly expressed isoform in melanocytes and melanoma (Goding & Arnheiter, 2019; Kawakami & Fisher, 2017). Previous genome-wide analyses of melanoma gene expression and MITF-binding regions revealed that MITF target genes are enriched in DNA replication, repair, and mitosis(Strub *et al*, 2011).

Recently, we have found that MITF silencing impairs the survival and proliferation of gastrointestinal stromal tumors (GIST)(Proaño-Pérez *et al*, 2023b). These tumors are the most frequent sarcomas and are mainly derived from interstitial cells of Cajal (ICC)(Casali *et al*, 2018), which are responsible for gastrointestinal tract motility. GIST are heterogeneous, and the most frequent drivers are active mutations that occur in *KIT* (60–70% of cases) and *PDGFRA* (10–15%) and are mutually exclusive (Blay *et al*, 2021). The major MITF isoform in GIST is MITF-A, the longer MITF isoform (Proaño-Pérez *et al*, 2023b). Information regarding MITF gene targets in GIST deserves further investigation. To gain insight into the role of MITF in GIST, we performed a ChIP-seq analysis to identify MITF binding sites and MITF-regulated genes in GIST. The study shows MITF involvement in an extensive set of genes mainly required for lysosome biogenesis, synaptic vesicles, and autophagy. Recent research highlights the critical role of autophagy, a catabolic process that can either promote cancer cell survival or death, in the behavior and treatment resistance of gastrointestinal stromal tumors (GISTs). Studies have shown that various molecular mechanisms promoting autophagy contribute to imatinib resistance (Sui *et al*, 2022; Gao *et al*, 2023). These findings indicate that targeting autophagy-related pathways might offer new therapeutic strategies. Autophagy intersects with extracellular vesicle secretion, sharing common regulatory mechanisms and components, such as the mTOR pathway (Salimi *et al*, 2020).

Our in vitro experiments after MITF-silencing confirm the involvement of MITF in autophagy and differential protein content of extracellular vesicles after proteomic analysis. These findings suggest that the MITF-autophagy-EVs axis may play a crucial role in tumor growth and resistance in GIST.

## RESULTS

### MITF targets analysis by ChIP-seq in GIST cells

A ChIP-seq was performed to explore which genomic regions of GIST are potential targets for MITF. The experiments were conducted in GIST-T1 and GIST48, harboring clinically representative KIT mutations. The total number of reads and normalized tags used for peak calling are shown in supplementary Tables I and II. Those results are derived from the raw results based on MacS (v2.1.0), which was filtered and kept the most significant peaks (Supplementary Material 1).

Known motifs were identified in GIST-T1 and GIST48 with the findMotifsGenome program of the HOMER package using default parameters and input sequences comprising +/- 100 bp from the center of the top 2500 peaks. The highest ranking motif 5’-GTCATGTGAC-3’ was found in GIST-T1 and GIST48 with p-values of 1e-554 and 1e-484, respectively, and coverage of 68.44% in GIST-T1 and 67.31% in GIST48. Predicted *de novo* motifs were found with p-values of 1e-633 and 1e-598, respectively, and coverage of 66.09% in GIST-T1 and 71.85 % in GIST48 (Figure 1A). A global ChIP-seq data analysis shows similar MITF-occupied loci and higher in proximal promoters, 5’ UTR (Figure 1B). Next, we compare peak regions in GIST-T1 and GIST48. A similar distribution of peak tag numbers was found in GIST-T1 and GIST48, with a Pearson correlation of 0.641 (Figure 1C-D). Tag distributions (using bigWIG metrics) in Merged Regions (= all peak regions; +/- 5 kb) are also shown (Figure 1E). Moreover, the Venn diagram shows common peaks (497) in both cell lines (Figure 1F), thus supporting a common role of MITF downstream KIT in a GIST-cell context.

**Figure 1.**
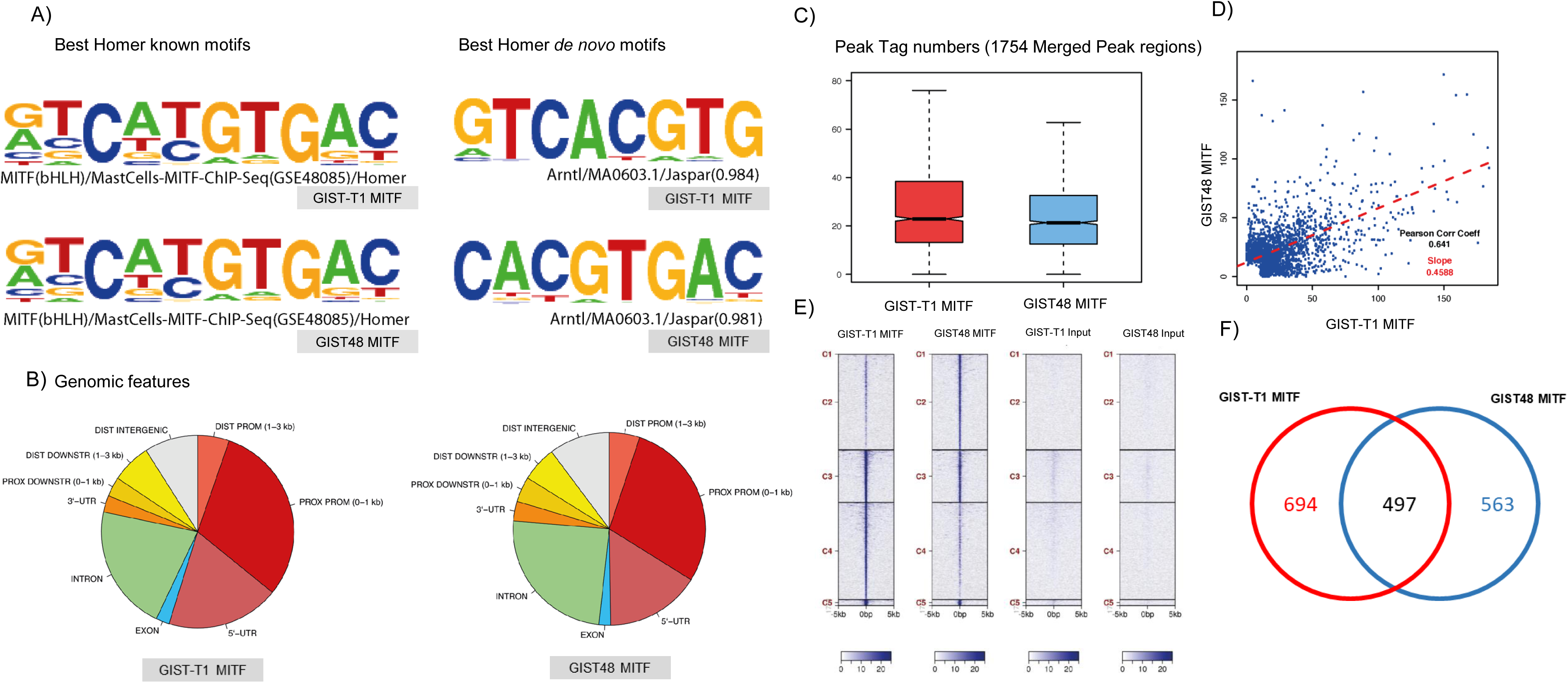
MITF enriched motifs and genomic features in target regions. ChIP-seq analysis. A) Left: Sequence logos showing the best de novo motifs annotated as enriched by the HOMER server for GIST-T1 MITF and GIST48 MITF samples with p-values of 1e-633 and 1e-598, respectively. These motifs cover 66.09% and 71.85% of the characterized targets. Right: Sequence logos showing the best-known motifs annotated as enriched by the HOMER server in GIST-T1 MITF and GIST48 MITF samples with p-values of 1e-554 and 1e-554, respectively. These motifs cover 64.44% and 67.31% of the characterized targets. B) Left: Pie chart of genomic features annotated for the GIST-T1 MITF sample. Right: Pie chart of genomic features annotated for the GIST48 MITF sample. C) Peak signal box plot comparing the distribution of peak tag numbers between the samples (merged peak regions). The boxed area represents the center two quartiles (notched line = median), while the whiskers show the top and bottom quartiles without outliers.D) Peak correlation scatter plot for the tag numbers of GIST-T1 MITF sample against GIST48 MITF sample for each merged region. The slope (linear model) measures the average ratio of tag numbers between the two samples. E) Heatmap for tag distributions (using bigWig metrics) across merged regions (all peak regions; +/- 5 kb). Tag distributions are clustered using the k-means algorithm, considering five clusters (indicated by C1-C5), and sorted by decreasing average values within each cluster. F) The Venn diagram shows the number of merged regions that are sample-specific, overlapping, and thus common to both samples.

The analysis of the peaks identified a total of 2.778 genes (Supplementary Material 2), with 907 genes being common in both cell lines. The pathway enrichment analysis (FDR <0.05) of the genes showed KEGG enrichment in processes associated with lysosomes, synaptic vesicle cycle, mTOR signaling, and autophagy (Figure 2A). The gene ontology (GO) biological function was predominant in protein location and transport, metabolic process, and autophagy regulation (Figure 2B).

**Figure 2.**
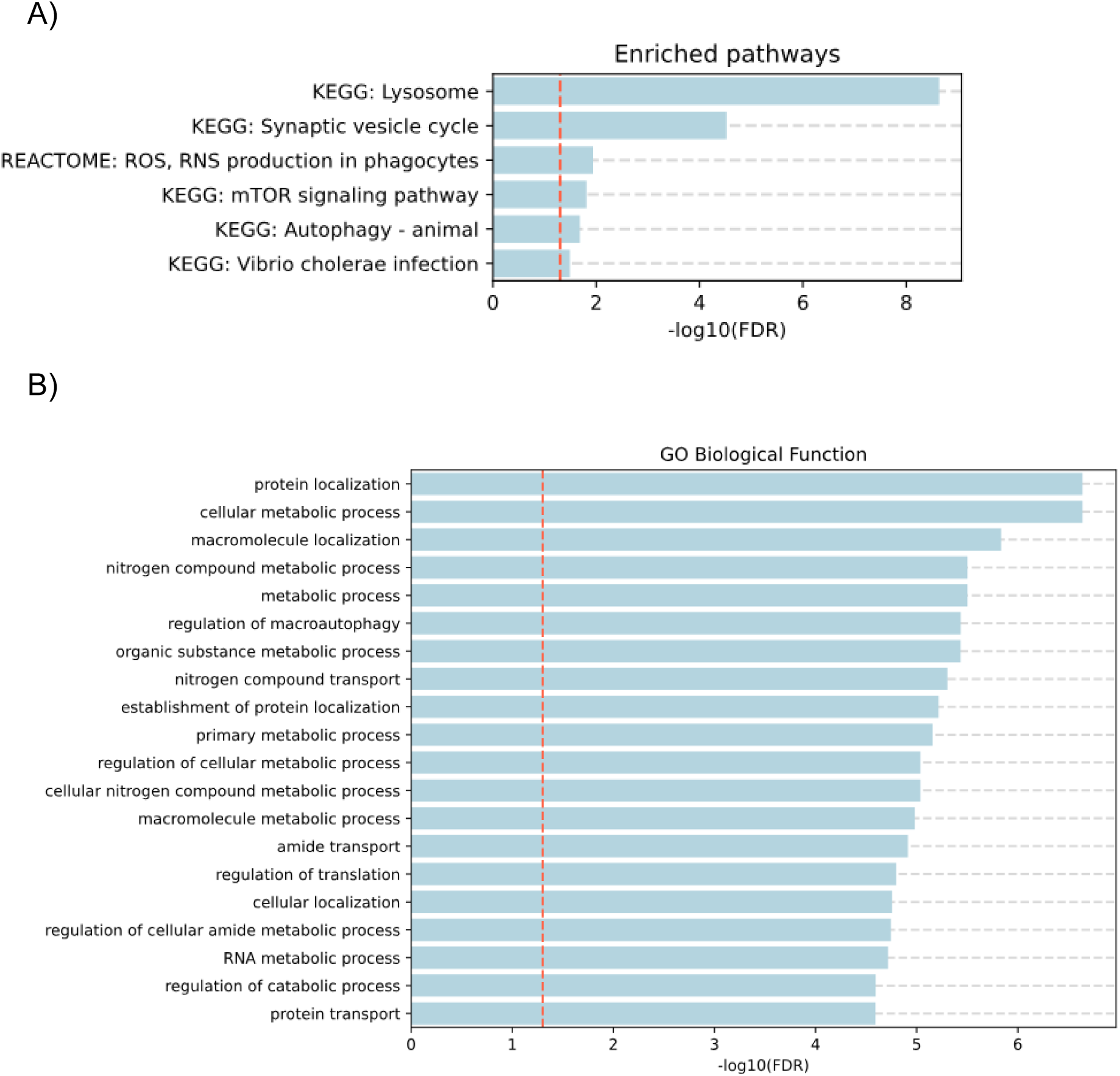
MITF-enriched pathways in GIST from ChIP-seq analysis data. A) Enriched pathways analysis and B) GO Biological function of common genes identified in the ChIP-seq study in GIST-T1 and GIST48.

### Identification of MITF-regulated genes in GIST cells

To identify potential target genes regulated by MITF, we used previously published shRNA sequences to silence MITF in the GIST-T1 cell line (Proaño-Pérez *et al*, 2023b). Following MITF silencing, RNA sequencing was performed. We analyzed the combined data against the non-targeting (NT) shRNA control and found 3.345 genes down-regulated and 2.196 genes up-regulated in MITF-silenced cells compared to the control. Among these, 136 down-regulated and 147 up-regulated genes overlapped with those identified in ChIP-seq experiments (Figure 3A and Supplementary Material 3). Gene Ontology (GO) showed Golgi and membrane components enrichment (Figure 3B), and metabolic pathway analyses revealed significant involvement of the mTOR and PI3K–Akt signaling pathways (Figure 3C), consistent with the ChIP-seq data. The number of differentially expressed genes closely matched the number of genes described as involved in these pathways, highlighting MITF’s critical role in regulating mTOR signaling and PI3K-AKT pathways in GIST (Table 1). Detailed information on differentially expressed genes related to enriched pathways is provided in Supplementary Material 4 and Supplementary Figure 1.

Differentially enriched Metabolic pathways in RNAseq.

Table 1. Differentially enriched Metabolic pathways in RNAseq
| Category | Pathway | numDEInCat | numInCat | pvalue | Adj. pvalue |
| --- | --- | --- | --- | --- | --- |
| map00040 | Pentose and glucuronate interconversions | 24 | 33 | 9,50E-06 | 0,00083427 |
| map00010 | Glycolysis and Gluconeogenesis | 31 | 47 | 1,77E-05 | 0,00083427 |
| map04150 | mTOR signaling pathway | 167 | 355 | 1,90E-05 | 0,00083427 |
| map04151 | PI3K-Akt signaling pathway | 165 | 353 | 3,31E-05 | 0,00109318 |
| map00053 | Ascorbate and aldarate metabolism | 17 | 23 | 0,00015173 | 0,00400578 |
| map00340 | Histidine metabolism | 18 | 26 | 0,00048152 | 0,01059352 |
| map00270 | Cysteine and methionine metabolism | 19 | 29 | 0,00076854 | 0,01265965 |
| map00500 | Starch and sucrose metabolism | 27 | 45 | 0,00082742 | 0,01265965 |
| map00100 | Steroid biosynthesis | 11 | 14 | 0,00086316 | 0,01265965 |

**Figure 3.**
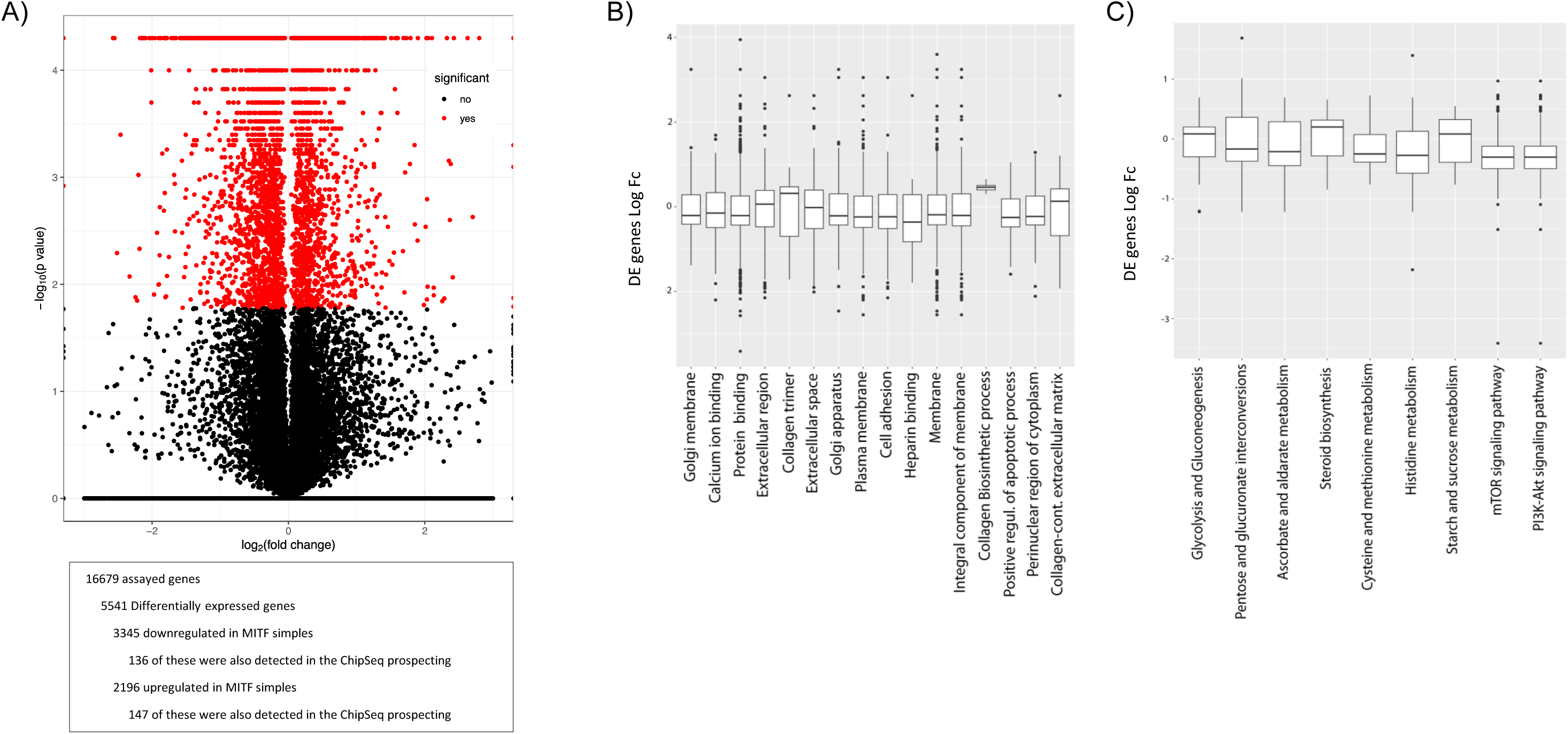
MITF silencing versus NT cells. RNA seq results. A) Volcano plot representing the log2 adjusted p values against the log2 fold change expression values obtained in the differential expression analysis. Red dots highlight the 5541 DE genes found as deregulated under an FDR < 0.05. B) Box plot analysis is based on the median, quartiles, maximum, minimum, and outlier values of the LogFCs of all transcripts contributing to the GO enrichment. LogFC values above zero correspond to the proportion of up-regulated genes contributing to the enrichment, and logFC values below zero correspond to down-regulated genes contributing to the enrichment. C) Same Box plot analysis based on all transcripts contributing to the enrichment of the metabolic pathways.

### MITF regulates genes required for autophagy in GIST cells

According to our GIST ChIP-seq results and RNA-seq data from MITF-silenced cells, which show differential expression of genes involved in the mTOR pathway and autophagy, we next conducted experiments to investigate the role of MITF in autophagy *in vitro*. We promoted autophagy using Torin 1, a mTORC1 inhibitor, or by cell starvation, both well-established methods for stimulating autophagy as outlined in autophagy guidelines articles (Ozturk *et al*, 2019; Klionsky *et al*, 2021). In both scenarios, we observed increased translocation of MITF to the nucleus (Figure 4A).

**Figure 4.**
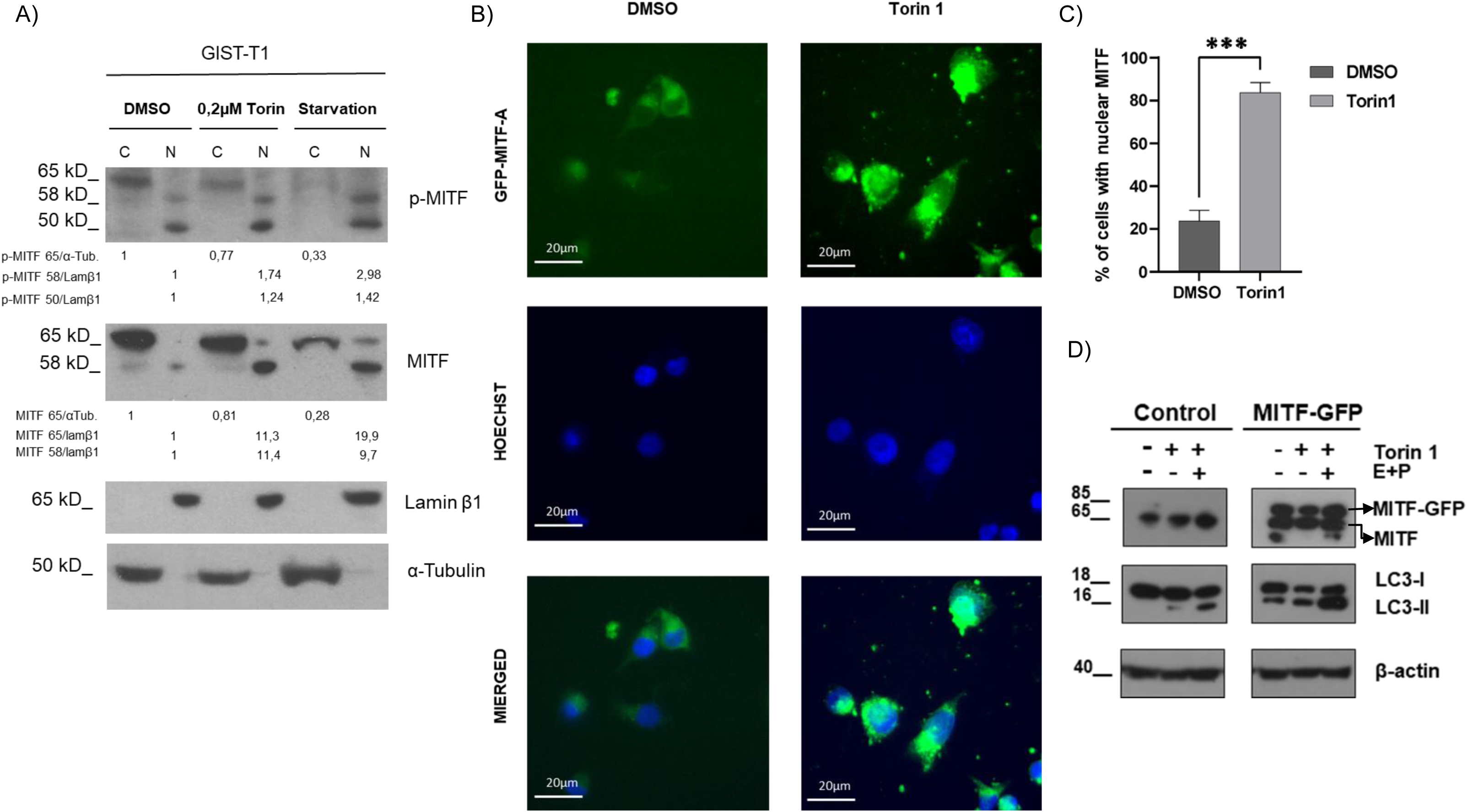
MITF is translocated to nuclei in autophagy, increasing LC3-II expression. A) MITF is translocated to nuclei of cells in autophagy. GIST-T1 cells were treated with Torin 1 (0,2µm) and starvation (EBBS solution) overnight. Cytoplasmic and nuclear fractions were isolated, and western blot for pMITF and MITF were performed; α-tubulin and Lamin β1 are markers for cytoplasm and nuclei, respectively, and were used to assess cellular fraction purification and as loading controls. B) GIST-T1 cells were transiently transfected with an MITF-GFP construct, incubated with DMSO or Torin1 (0,2µm), and analyzed by immunofluorescence microscopy. Hoechst dye was used to stain the nuclei (blue). C) Quantitative analysis of MITF nuclear translocation in the experimental setup shown in B (*** p<0.001; Unpaired T-test; mean ±SEM). D) Overexpression MITF-GFP leads to an increase in LC3II in autophagy induction. Western blot showing MITF overexpression in GIST-T1 cells and LC3I-II levels were assessed in Torin 1-treated or untreated cells. E64D (10 µg/ml) + pepstatin A (10 µg/ml) were used as lysosomal protease inhibitors. Beta-actin was used as a loading control.

To analyze MITF’s contribution to autophagy in GIST, we overexpressed MITF-GFP and performed an LC3II shift assay. LC3II serves as a marker for both growing and completed autophagosomes. Under control conditions, MITF-GFP expression was primarily cytoplasmic. Following Torin 1 treatment, there was an increase in MITF-GFP nuclear translocation and LC3II accumulation compared to control-treated cells (Figure 4B-D). To confirm that MITF increased autophagosome formation and that LC3II accumulation was not due to a block in autophagosome-lysosome fusion, we used lysosomal protease inhibitors E64D and pepstatin A. The LC3II levels remained higher in MITF-overexpressing cells, indicating that MITF promotes autophagosome formation (Figure 4D, E+P lanes).

To determine whether MITF was a rate-limiting factor in autophagy in GIST, we silenced MITF and assessed autophagy using GFP-LC3 expression. Cells undergoing autophagy can be identified by visualizing fluorescently labeled LC3 puncta. Our results showed that MITF silencing significantly reduced GFP-LC3 puncta levels, even under basal conditions (Figure 5A-B). Given that GFP-LC3 aggregates have been reported in autophagy-deficient cells and this approach has been questioned (Runwal *et al*, 2019), we further analyzed endogenous LC3II expression in GIST cells transduced with NT and MITF shRNAs, in the presence or absence of lysosomal inhibitors. MITF-silenced cells exhibited reduced endogenous LC3II levels (Figure 5C), which decreased proportionally with MITF levels.

**Figure 5.**
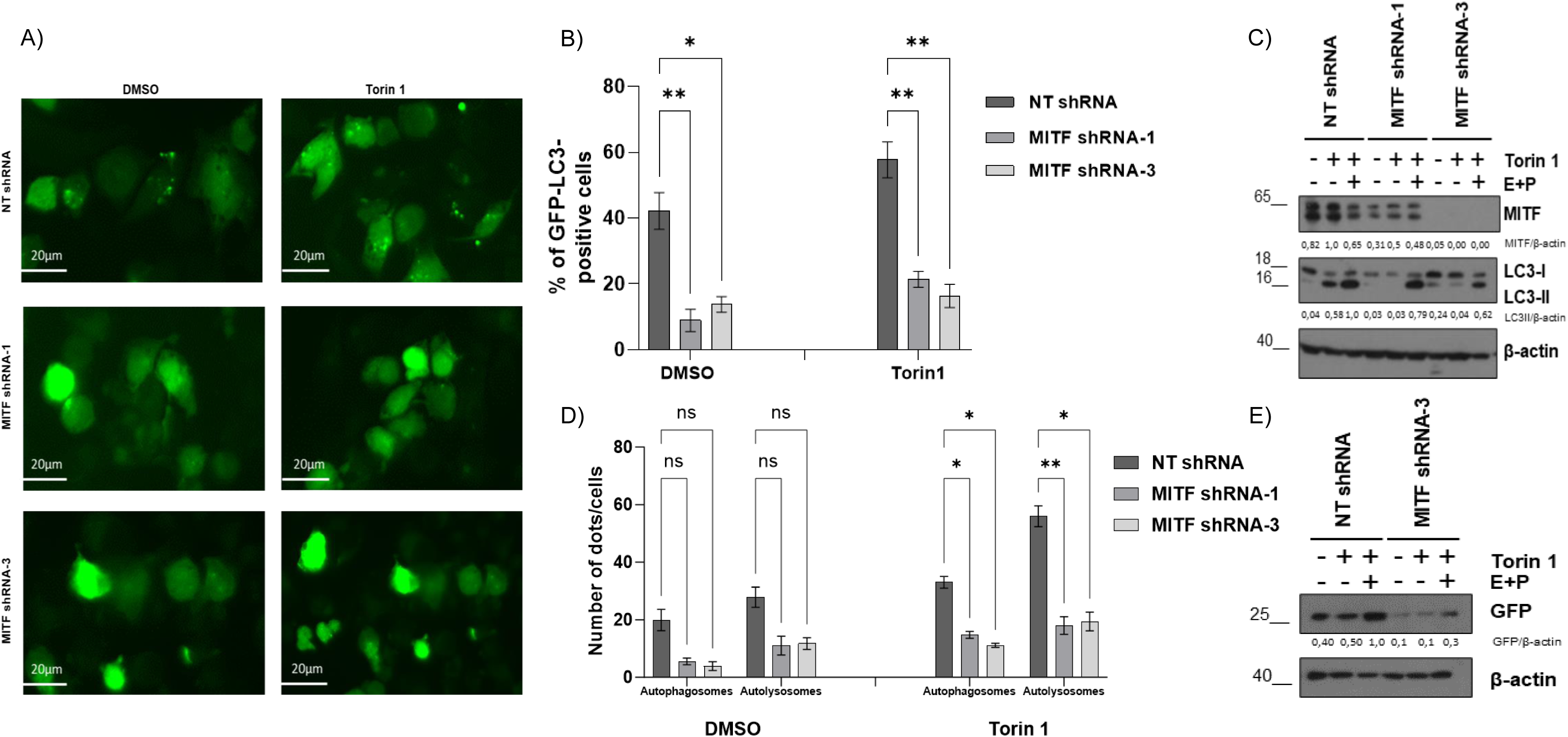
Autophagy is Diminished in MITF Silencing Cells after Torin1 Treatment. A) GFP-LC3 puncta formation was assessed by immunofluorescence microscopy in MITF-silenced GIST-T1 cells and their non-target control. B) Quantitative analysis of GFP-LC3 puncta formation as described in panel A (*p < 0.05, **p < 0.01; one-way ANOVA with Bonferroni’s post hoc test). C) LC3II expression is impaired in MITF-silenced cells. Western blot analysis was performed in cells transduced with non-target shRNA, MITF shRNA-1, and MITF shRNA-3 after autophagic induction with Torin1 and treatment with lysosomal protease inhibitors (E64D and pepstatin). The graph shows ratios of LC3II protein expression. Beta-actin serves as a loading control. D) Quantitative analysis of autophagosomes and autolysosomes (*p < 0.05; one-way ANOVA with Bonferroni’s post hoc test). E) GFP-LC3 lysosomal delivery and proteolysis assay. GIST-T1 cells transduced with MITF shRNA were transfected with GFP-LC3 plasmid and treated with DMSO or Torin1 plus E64D + pepstatin. Free GFP levels are shown in the blots. Beta-actin was used as a loading control.

Additionally, we performed an autophagy flux analysis in MITF-silenced cells. To assess autophagy flux, we used GIST-T1 stable cells expressing RFP-GFP LC3. This construct allows the measurement of autophagosomes and autolysosomes, as the GFP signal is quenched in lysosomes while the RFP signal is retained, resulting in the identification of autolysosomes(Kimura *et al*, 2007). After Torin 1 treatment, control cells showed increased autophagosomes and autolysosomes. Interestingly, these numbers decreased in MITF-silenced cells (Figure 5D and Supplementary Figure 2).

To further corroborate MITF’s involvement in autophagy flux in GIST, we tested GFP-LC3 lysosomal delivery and proteolysis. In this process, GFP-LC3 is delivered to lysosomes, where LC3 is degraded while GFP remains resistant to hydrolysis. Free GFP expression can thus be used to monitor the breakdown of autophagosomal cargo. Our data showed an accumulation of free GFP in cells treated with Torin 1, indicating increased autophagy flux. However, this accumulation was reduced in MITF-silenced cells. These results confirm that MITF regulates autophagy in GIST, which aligns with ChIP-seq and transcriptomic results (Figure 5E).

### MITF silencing does not affect the size and number of extracellular vesicles (EVs) released in GIST cells

MITF ChIP-seq analysis revealed that MITF may be involved in lysosome biogenesis, vesicle cycle, and protein transport (Figure 2). As part of the vesicular communication system, EVs and autophagy work together to support cellular adaptation and intercellular communication. This collaboration facilitates the selection and secretion of damaged cellular material, a process orchestrated by the endoplasmic reticulum, the Golgi apparatus, and lysosomes (Jahangiri *et al*, 2022).

Hence, we aimed to investigate whether MITF influences the biogenesis of EVs or their protein cargo in GIST. EVs released by GIST cells facilitate intercellular communication, promoting tumor growth and cancer progression (Atay *et al*, 2014). We isolated EV from NT shRNA, MITF shRNA-1, and MITF shRNA-3 cells using size exclusion chromatography (SEC).

The presence of tetraspanins CD9, CD63, and CD81, identified by flow cytometry, was used to confirm the EV-enriched fractions after SEC isolation. Our results showed similar flow cytometry profiles in control and MITF-silenced cells (Figure 6A-B) with no significant differences in any marker assessed. Nanoparticle tracking analysis (NTA) was further used to determine EV concentration and size distribution. EV concentration yielded around 4-6×10^9^ part/ml in all three conditions, indicating that MITF silencing did not affect the number of EVs released (Figure 6C).

**Figure 6.**
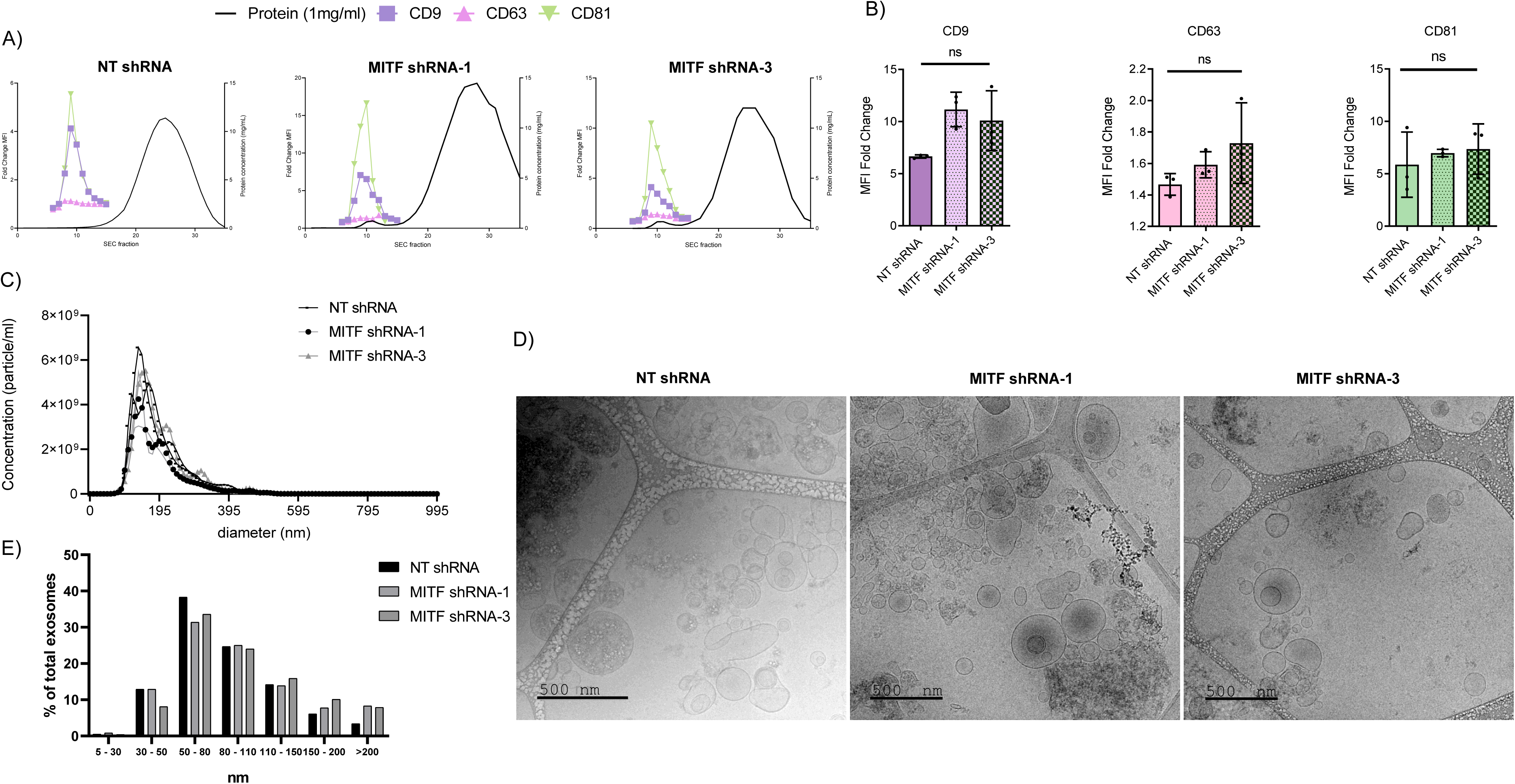
Characterization of Enriched Fractions of EVs from GIST MITF shRNA-1, MITF shRNA-3, and NT shRNA. A) Flow cytometry characterization of EVs in GIST. Enriched fractions of EVs from GIST MITF shRNA-1, MITF shRNA-3, and NT-shRNA using EVs markers: CD9, CD63, and CD81. B) Fold change in mean fluorescence intensity (MFI) correlated with protein concentration (1mg/ml). C) Nanoparticle Tracking Analysis (NTA) measuring EVs’ diameter and concentration. D) Characterization of the size and morphology of EVs by cryo-TEM TEM (cryo-transmission electron microscopy). Scale bars represent 500 nm. E) Percentage of EVs analyzed from cryo-TEM samples.

Furthermore, isolated vesicles were analyzed using cryo-electron microscopy. Measurements from cryo-electron microscopy images confirmed the NTA analysis results, showing a significant peak of abundant vesicles around 50-150 nm (Figure 6D-E). Our findings conclude that MITF silencing does not affect the size and number of EVs in GIST-T1 cells.

### Proteomic analysis of EVs. Silencing MITF alters EV content related to ER stress and cytokinesis

Next, we performed a proteomic analysis of the isolated EVs from MITF-silenced cells. The complete set of quantified proteins is provided in Supplementary Material 5 (SM5_1). To gain insights into the potentially altered signaling pathways in these EVs, we conducted a Gene Set Enrichment Analysis (GSEA) comparing MITF-silenced cells with control cells (Figure 7A and Supplementary Material 5 (SM5_2 and SM5_3)). The results indicated that MITF-silenced cells were mainly enriched in pathways related to the cytoskeleton, cytokinesis, glycoprotein metabolic processes, and protein folding and maturation in the endoplasmic reticulum (ER). Signaling pathways involved in PI3K and MAPK kinases, ER processing, and lysosomes were also enriched.

**Figure 7.**
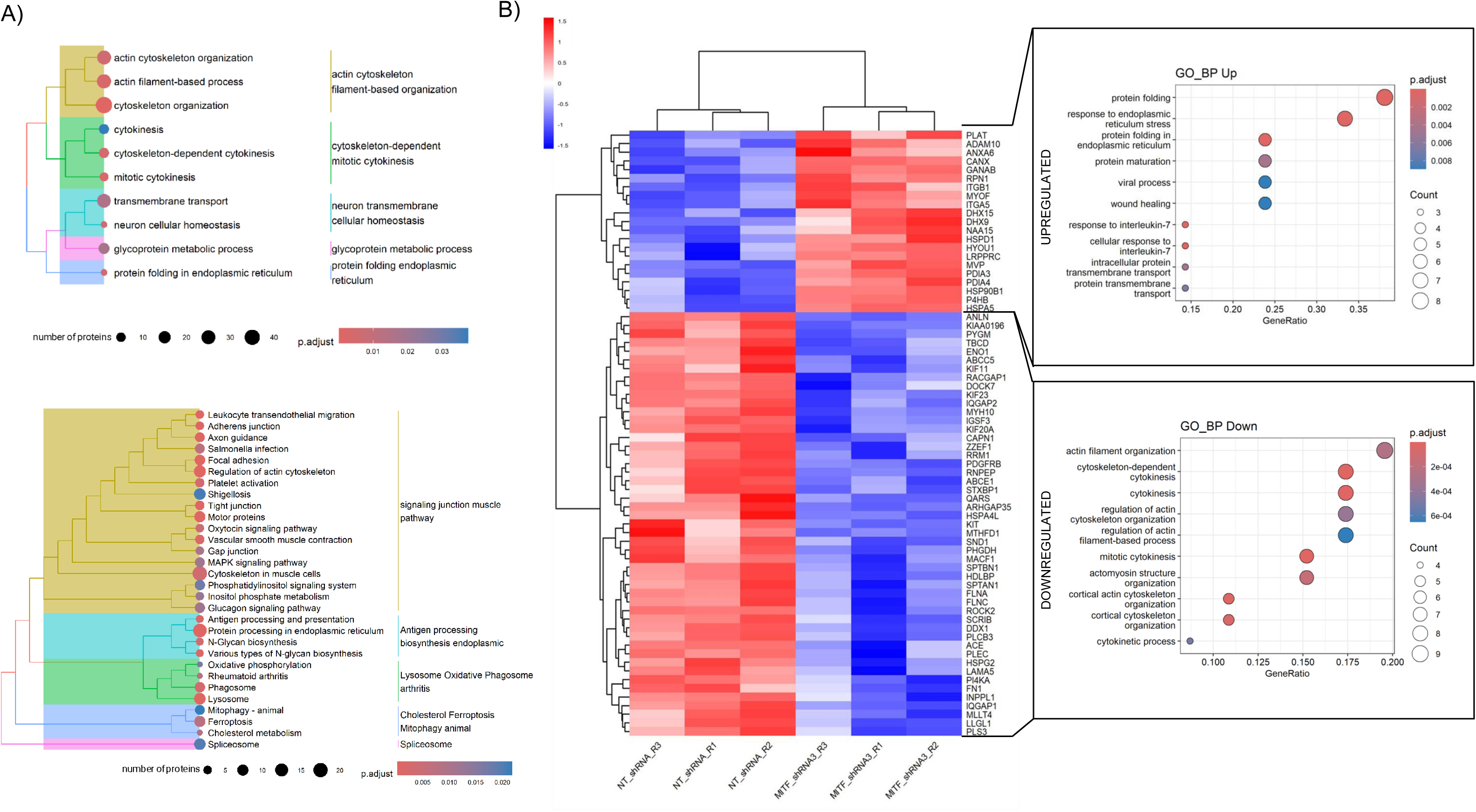
Proteomic Analysis of Exosomes Derived from MITF Silenced Cells Compared to NT Cells. A) Gene Set Enrichment Analysis (GSEA) of all proteins present in exosomes using Gene Ontology Biological Process (GO BP) terms (upper panel) and KEGG terms (lower panel). Circle size indicates the number of associated proteins, and numbers represent specific numerical values. B) Heatmap displaying differential gene expression for proteins between NT-shRNA control cells and MITF-silenced cells (MITF_shRNA3) (left panel), and Over-Representation Analysis (ORA) of up-regulated and downregulated proteins (right panel). Red and blue shadings in the heatmap indicate higher and lower relative expression levels, respectively.

For a more focused, stringent analysis, we filtered the proteins to include only those represented by at least 25 peptides, with a q-value <0.05 and a logFC of ±0.5849. As shown in the heatmap in Figure 7B (left panel), 49 proteins were downregulated in the EVs from MITF-silenced cells, while 21 proteins were up-regulated. Using this list of differentially expressed proteins, we performed an Over-Representation Analysis (ORA) to identify enriched biological functions (Figure 7B, right panel, and Supplementary Material 5 (SM5_4 and SM5_5)).

The ORA analysis revealed that biological functions related to protein folding and ER stress were up-regulated in the MITF-silenced EVs. In contrast, functions associated with cytokinesis and cell shape (including cytoskeleton and actin organization) were enriched among the downregulated proteins. These findings align with observations that MITF silencing negatively impacts cell division and survival, reinforcing previously reported data(Proaño-Pérez *et al*, 2023b).

### MITF silencing impairs KIT content in EVs in GIST

Proteomic data from EVs showed that KIT content was reduced in MITF silencing. The presence of KIT in exosomes has been reported to increase GIST cell invasion (Atay *et al*, 2014), and the protein levels have therapeutic value (Atay *et al*, 2018). Next, we validated these results by western blot and characterized the Evs biochemically using various biomarkers: Alix, Calnexin, TSG101, Flotilin, Syntenin, CD9, and Cox following MISEV guidelines (Welsh *et al*, 2024). Our findings revealed a significant reduction in KIT levels within EVs following MITF silencing (Figure 8).

**Figure 8.**
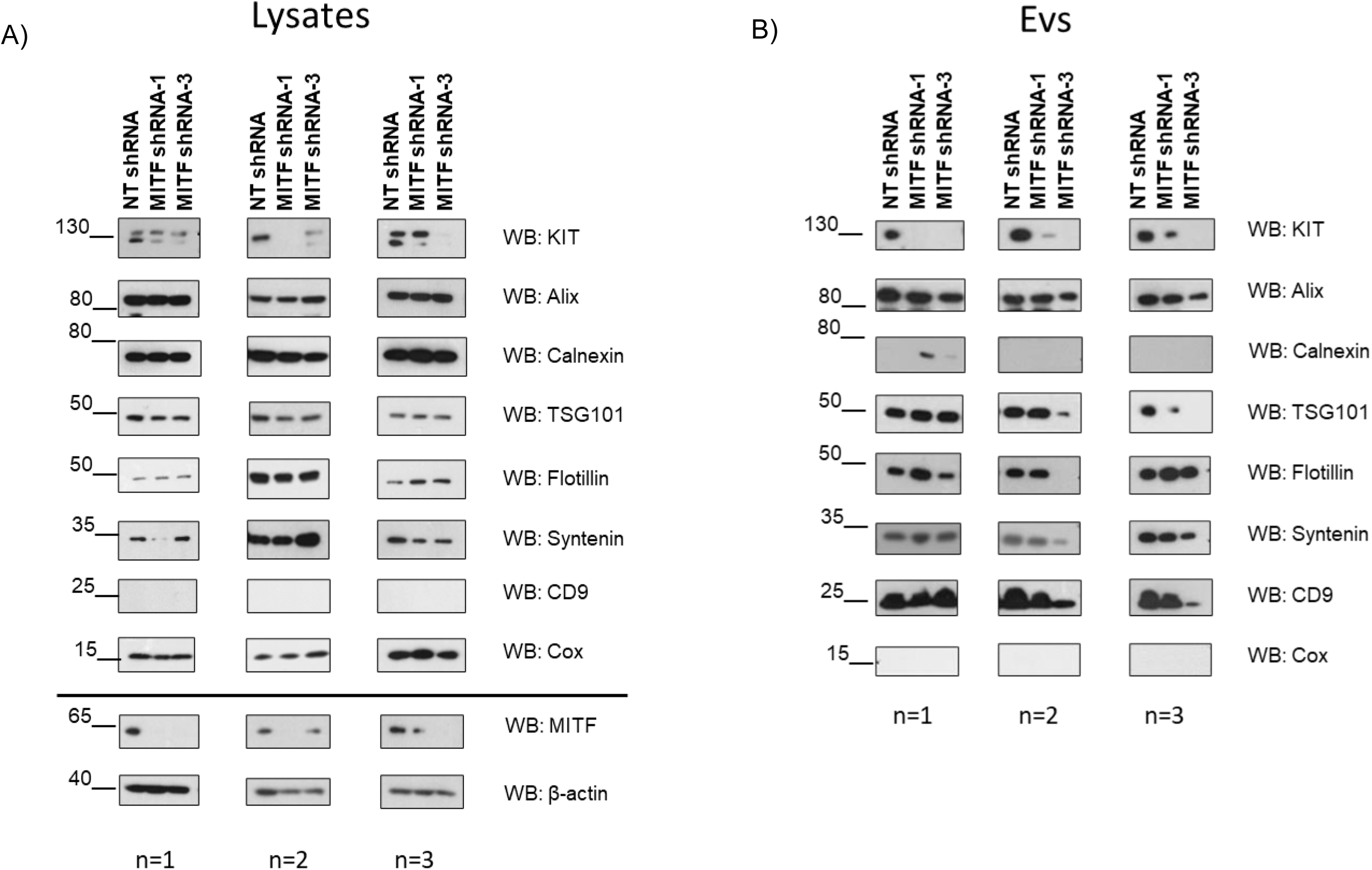
KIT protein is Reduced in Lysates and Exosomal Content in MITF-silenced cells. Cells transduced with Non-target shRNA, MITF shRNA-1, or MITF shRNA-3 were analyzed. Lysates from GITS-T1 and exosomal content were analyzed by Western blot. Exosome markers (CD69, syntenin, flotillin, Alix, and TSG101), mitochondrial cytochrome-c oxidase (COX), endoplasmic reticulum control (Calnexin), KIT, MITF, and β-actin were examined by Western blot. N=3 independent experiments

Altogether, our results suggest that MITF plays a crucial role in modulating the protein cargo of EVs, potentially influencing the invasive capabilities and progression of GIST through altered intercellular communication mediated by KIT-containing exosomes.

## DISCUSSION

MITF has garnered significant attention for its multifaceted cellular physiology role, particularly in melanoma. However, emerging evidence suggests that MITF may also be pivotal in the pathology of GIST(Proaño-Pérez *et al*, 2023b). This study delves into MITF’s role in GIST by leveraging ChIP-seq and transcriptomics.

MITF ChIP-seq in melanoma (Human 501 Mel cells) identified targets involved in DNA replication, repair, and mitosis (Strub *et al*, 2011). Our study reveals that prominent targets of MITF in GIST include genes involved in lysosome biogenesis, mTOR signaling, and autophagy. Lysosomes are essential for cellular metabolism and play a critical role in autophagy, a catabolic process necessary for cell homeostasis with physiological and pathophysiological roles (Mizushima, 2007). This pathway is typically activated under stress conditions, such as starvation or nutrient amino acid deprivation (Gozuacik & Kimchi, 2004). In cancer, autophagy serves a dual role: it acts as a tumor-suppressive mechanism in the early stages by preventing tumor initiation and progression, but in later stages, it supports tumor cell survival and growth (Singh *et al*, 2018). The upregulation of autophagy has been reported in nearly all cancers and may promote metastasis (Ma *et al*, 2011).

Our study found that MITF overexpression increases LC3II, an autophagy marker, while MITF silencing reduces LC3II levels and decreases autophagosome and autolysosome formation. These findings indicate that MITF is involved in autophagy and autophagic flux in GIST. MITF’s role in autophagy has also been observed in melanoma (SK-MEL-28 cells) and HeLa cells (Ozturk *et al*, 2019). Additionally, TFEB and TFE3, members of the MITF/TFE family of bHLH-Zip transcription factors, are also related to autophagy (Settembre *et al*, 2013; Martina *et al*, 2014; Theodosakis *et al*, 2022). The inhibition of autophagy through the DREAM complex, by depleting the DREAM regulatory kinase DYRK1A or its target LIN52, enhances imatinib-induced cell death in GIST(Boichuk *et al*, 2013). DYRK1A appears in our ChIP-seq data, and transcriptomic data shows a significant reduction of DYRK1A after MITF silencing. MITF-silenced transcriptomic data shows notable enrichment of the mTOR and PI3K-Akt signaling pathways. The consistent deregulation of genes within these pathways underscores MITF’s significant influence. Intriguingly, MITF silencing results in a decrease in signaling associated with mTOR and PI3K. Interestingly, PI3K/AKT/mTOR pathway activation has been associated with secondary imatinib (IM) resistance in patients with GISTs (Li *et al*, 2015). Autophagy has also been correlated with IM resistance. Rubin and colleagues demonstrated that IM induces autophagy as a survival pathway in quiescent GIST cells (Gupta *et al*, 2010). Moreover, in a PAK-dependent manner, autophagy-related protein 5 (ATG5) enhances autophagy and promotes IM resistance in GIST (Gao *et al*, 2023). Thus, regulators of autophagy have potential therapeutic applications (Li *et al*, 2020). Additionally, It has been proposed that IM alters the metabolic phenotype of GIST through ROS and HIF-1α, contributing to IM resistance in vivo and in vitro (Xu *et al*, 2020). Our transcriptomic data shows significant reductions in PAK1 and HIF-1α in MITF-silenced cells. Investigating MITF and its associated targets could lead to developing targeted therapies that inhibit drug resistance pathways.

Differentially expressed genes in our transcriptomic analysis are involved in the transport and delivery of proteins and the maturation of early endosomes. Recent evidence highlights exosomes as endosome-derived small EVs that participate in cell-to-cell communication and the delivery of proteins and RNA (Raposo & Stoorvogel, 2013). Importantly, EVs are promising novel biomarkers for clinical diagnosis (Lin *et al*, 2015). Moreover, many carcinomas rely on establishing complex communication networks within and between the stroma, where EVs may play a crucial role (Löffek *et al*, 2006).

Autophagy has been linked to EV production and vice versa, playing a role in cellular adaptation and communication. This interplay can occur at different levels: autophagy stimulation may increase or decrease EV secretion, contributing to either the progression or suppression of diseases (Xing *et al*, 2021). Indeed, the cargo content of EVs may play an important role in regulating autophagy in recipient cells (Salimi *et al*, 2020). Cell signaling and stress conditions can affect the interplay between EVs and autophagy (Hessvik *et al*, 2016). The contents of EVs vary based on their mode of biogenesis, cell type, and physiological conditions (Abels & Breakefield, 2016). EVs typically exhibit proteins associated with biogenesis, such as ALIX and TSG101, and proteins involved in EV formation and release, such as RAB27A, RAB11B, and ARF6. Additionally, EVs contain various tetraspanins, including CD63, CD81, and CD9, and proteins involved in signal transduction (e.g., EGFR), antigen presentation (MHC I and MHC II), and other transmembrane proteins (e.g., LAMP1, TfR). Notably, proteins associated with the ER, Golgi, and nucleus are not found in EVs(Théry *et al*, 2002).

Our study demonstrates that while EVs’ number and size are unaffected by MITF silencing, their cargo exhibits a distinct differential signature. Enhanced responses of ER stress proteins and signals related to protein folding and maturation were observed. The unfolded protein response (UPR) typically promotes cell survival, but unresolved ER stress can lead to apoptosis (Ron & Walter, 2007). ER stress may trigger either pro-survival or pro-apoptotic responses depending on the severity and duration of the stress stimuli, which in turn affects the differential activation of the UPR (Rutkowski *et al*, 2006).

Additionally, the EV cargo shows downregulation of proteins involved in cytokinesis and cell division, and this aligns with previous data indicating that MITF silencing reduces cell viability, increases apoptosis, and arrests the cell cycle (Proaño-Pérez *et al*, 2023b). Our transcriptomic analysis reveals a decrease in the expression of the anti-apoptotic gene BCL2 and the cyclin-dependent kinase CDK2, which is crucial for the S-G2/M phase transition, further reinforcing our previous findings (Proaño-Pérez *et al*, 2023b).

EVs may serve as immune system suppressors, modulators of tumor promotion, and biomarkers for several cancers (Graner *et al*, 2018). Previous reports have shown that exosome content from melanoma cells promotes metastasis in bone marrow progenitor cells(Peinado *et al*, 2012), indicating that EV content can influence the conversion of normal cells into oncogenic cells. Similarly, the accumulation of circulating exosomes is enhanced in the peripheral blood of patients with metastatic GIST compared to those with primary disease GIST. Proteomic analysis revealed that increased cell metastasis depends on oncogenic KIT transported in these vesicles (Atay *et al*, 2014). Furthermore, circulating levels of KIT+ exosomes in GIST patients correlate with tumor burden (Atay *et al*, 2014, 2018).

More recently, systemic mastocytosis (SM)-derived EVs have been shown to activate hepatic stellate cells, leading to the development of liver disease through functionally active KIT from SM-EVs, highlighting aspects of liver pathology in mastocytosis (Kim *et al*, 2018). Interestingly, MITF impairment reduces KIT expression in both GIST and HMC-1, a cellular model of mastocytosis (Proaño-Pérez *et al*, 2023b, 2023a). In GIST, MITF silencing resulted in reduced KIT protein expression ^15,75^. In this study, we consistently observed decreased KIT protein levels in EVs from MITF-silenced GIST-T1 cells, which may attenuate the KIT-dependent paracrine activity of EVs.

This study underscores MITF as a promising target for therapeutic intervention in GISTs due to its regulatory influence on crucial pathways involved in tumor growth, autophagy, and metastasis. Further exploration of MITF’s molecular mechanisms and its interaction with downstream effectors could provide valuable insights into GIST pathobiology and guide the development of novel therapeutic strategies targeting MITF.

## MATERIAL AND METHODS

### Cells, antibodies, and reagents

Human GIST cell lines were kindly provided by Dr. S. Bauer (University Duisburg-Essen, Medical School, Essen, Germany). Imatinib-resistant GIST48 (mutations p.Asp820Ala; p.Val560Asp) (RRID:CVCL_7041) (Bauer *et al*, 2006) cells were maintained in Ham’s F-10 media (Lonza) supplemented with 15% FBS, 1% L-glutamine, 50 units/ml penicillin, and streptomycin, 30 mg/ml bovine pituitary extract and 0.5% MITO+ Serum Extender (Fischer Scientific, Pittsburg, PA). Imatinib-sensitive GIST-T1 (mutation KIT; Simple; p.Val560_Tyr578del) (RRID:CVCL_4976) has been published previously (Serrano *et al*, 2019). The Mycoplasma test is performed routinely in all cell lines used. Mouse anti-KIT (clone Ab81), anti-CD81 (cloneG0709), and rabbit anti-GFP were purchased from Santa Cruz Biotechnology, Inc. (Santa Cruz, CA, USA). Anti-MITF (clone D5G7V), anti-Alix, anti-CD9 (clone 13174), and anti-LC3 I-LC3II (clone 81631) were obtained from Cell Signaling Technology, Inc (Danvers, MA, USA). Mouse anti-COX IV (A-21347) was from Invitrogen. Mouse anti-β-actin (clone AC-40) was from Sigma (St. Louis, MO, USA). Anti-syntenin (ab8133267), anti-TSG101 (ab30871), anti-CD63 (ab8219), and anti-calnexin (ab22595) were purchased from Abcam technology (Abcam, Cambridge, UK). Anti-mouse and anti-rabbit IgG peroxidase Abs were acquired from DAKO (Carpinteria, CA, USA) and Biorad (Hercules, CA, USA), respectively. Flotillin1 anti-mouse (#610820) was obtained from BDBiosciencies.

### Chromatin Immunoprecipitation Sequencing (ChIP-seq)

GIST-T1 and GIST48 samples were frozen at -120°C and sent to Active Motif (Carlsbad, CA) for ChIP-seq (RRID:SCR_001237). When samples arrived, cells were fixed with 1% formaldehyde for 15 min and quenched with 0.125 M glycine. In addition, chromatin was isolated by adding lysis buffer, followed by disruption with a Dounce homogenizer. Moreover, lysates were sonicated, and the DNA sheared to an average length of 300-500 bp with Active Motif’s EpiShear probe sonicator (cat# 53051). Besides, the genetic DNA (Input) was prepared by treating chromatin with RNase, proteinase K, and heating for cross-linking. Moreover, the process was followed by SPRI beads cleanup (Beckman Coulter) and quantitation through Clariostar (BMG Labtech). Finally, extrapolating the original chromatin volume allowed the determination of the total chromatin yield.

An aliquot of chromatin (30 µgs) was precleared with protein G agarose beads (Invitrogen). Genomic DNA regions of interest were isolated using 5 µg of antibody against MITF (Active Motif, cat# 39789, lot# 11313002). Complexes were washed, eluted from the beads with SDS buffer, and subjected to RNase and proteinase K treatment. Cross-links were reversed by incubation overnight at 65 °C, and ChIP DNA was purified by phenol-chloroform extraction and ethanol precipitation. Quantitative PCR (qPCR) reactions were carried out in triplicate on specific genomic regions using SYBR Green Supermix (Bio-Rad). The resulting signals were normalized for primer efficiency by carrying out qPCR for each primer pair through Input DNA. Illumina sequencing libraries were prepared from the ChIP and Input DNAs by the standard consecutive enzymatic steps of end-polishing, dA-addition, and adaptor ligation. This process was performed in an automated system (Apollo 342, Wafergen Biosystems/Takara). Finally, after the last PCR amplification step, the resulting DNA libraries were quantified and sequenced on Illumina’s NextSeq 500 (75 nt reads, single end).

### Bioinformatic analysis CHIP-seq

Fastq files were mapped on the human genome hg38 version with BWA (RRID:SCR_010910) (Li & Durbin, 2010) using (aln/samse) algorithm and default settings. Only reads that passed the purity filter of Illumina were aligned with no more than two mismatches and mapped uniquely to the genome mapping quality ≥ 25 were used in the subsequent analysis. Moreover, duplicate reads were removed.

Alignments were extended *in silico* at their 3’-ends to 200 bp average genomic fragment length in the size-selected library and assigned to 32-nt binds along the genome. The resulting histograms (genomic signal maps) were stored in bigWig files. Peaks were called using MACS (v2.1.0) (Zhang *et al*, 2008) with a cutoff of p-value = 1e-7. Peaks ENCODE in the blacklist or false ChIP-seq peaks were removed. Active Motifs’ proprietary analysis program used signal maps and peak locations as input data.

For each sample, the enrichment of motifs was assessed and graphically represented using HOMER. [http://homer.ucsd.edu/homer/] (RRID:SCR_010881) considering regions of 200 bp surrounding the summit of the top 2,500 peaks (based on MACS2 p-values).

Finally, annotations of enrichments in gene ontologies GO, metabolic KEGG (RRID:SCR_012773), and UniProt keywords were obtained from the STRING (RRID:SCR_005223) Server V12 web server (Szklarczyk *et al*, 2023).

### Transcriptomic analysis

Total RNAs from GIST-T1 transduced with MITF shRNA -1, MITF shRNA -3, or NT shRNA, as a control, were obtained in triplicates. Samples were sent at -80°C under strict conditions to MACROGEN for RNA-seq analysis. Firstly, a DNAse kit was used to prevent DNA contamination. Second, an appropriate setup for the library prep process was selected depending on the types of RNA. In addition, poly-A tail purified mRNA was obtained using an mRNA purification kit; for non-coding RNAs (LINC-RNA), a ribo-zero RNA-removal Kit was used to purify RNA of interest. Fourth, the random fragment was purified of RNA for short-read sequencing. The fifth step was to perform an RT-PCR to convert fragmented RNA into cDNA. As a sixth step, ligated adapters were inserted at both ends of the cDNA fragments. In the seventh step, select pieces whose insert sizes are between 200 and 400 bases after PCR amplifying cDNA fragments. In the final step, paired-end sequences from both ends of the cDNA were sequenced based on the read length.

### Bioinformatic analyses RNA seq

The analysis method was as follows: Overall reads’ quality, total bases, total reads, GC (%), and basic statistics were calculated. Low-quality reads adapter sequences, contaminant DNA, or PCR duplicates were removed to reduce biases in the analysis. Trimmed reads were mapped to reference genome with HISAT2 (RRID:SCR_015530), a splice-aware aligner. Transcripts were assembled by StringTie with aligned reads. Expression profiles were represented as read count and normalization values, which were calculated based on transcript length and depth of coverage. Normalization values were provided as FPKM (Fragments Per Kilobase of transcript per Million Mapped reads) / RPKM (Reads Per Kilobase of transcript per Million mapped reads) and TPM (Transcripts Per Kilobase Million).

#### Preprocessing

Quality control on Fastq libraries was performed using FastQC (RRID:SCR_014583) (Andrews & others, 2019). Subsequently, Fastq files were preprocessed using Cutadapt 1.18 (Martin, 2011) and Prinseq-lite-020.4 (Schmieder & Edwards, 2011) to eliminate primers and low-quality sequences.

#### Mapping and differential expression analyses

Preprocessed Fastq files were mapped on the human genome GRCh38 version provided by the Ensembl release 108 (Martin *et al*, 2023), using TopHat 2.1.1 (RRID:SCR_013035) (Kim *et al*, 2013). The overall read mapping rates successfully ranged from 87 to 92% of reads mapped. Bam files resulting from mapping with TopHat were used as input to Cufflinks/Cuffdiff (RRID:SCR_014597/ RRID:SCR_001647) to perform a differential expression (DE) analysis between the MITF-silenced study group (constituted 6 samples) and the control group (constituted 3 samples) as described elsewhere (Trapnell *et al*, 2012). Differentially expressed genes supported by a Benjamini-Hochberg False Discovery Rate (FDR) correction < 0.05 were considered statistically significant. Statistical and graphical analyses were performed using CummeRbund (RRID:SCR_014568) (Goff *et al*, 2013).

All differentially expressed genes were provided with annotations (regarding gene names, descriptions, gene ontologies (GO), enzyme commission (ECs) numbers, and metabolic pathways) using Biomart (RRID:SCR_019214) and Ensembl (RRID:SCR_002344) (Martin *et al*, 2023) and the KEEG (Repana *et al*, 2019)knowledge database. Finally, an enrichment analysis of GOs and metabolic pathways was performed using GOseq (Young *et al*, 2010) based on the differential expression results. To perform these analyses, all assayed genes were previously provided with annotations of GOs and metabolic pathways retrieved from Ensembl and Biomart (Martin *et al*, 2023; Rainer *et al*, 2019) and the KEGG database (Repana *et al*, 2019), respectively. Enriched GO terms and metabolic pathways supported by p-value corrected by FDR <

0.05 in the resulting Wallenius and/or Sampling distribution were considered statistically significant. The above-described steps for RNAseq data processing, differential expression, and enrichment analysis were executed using the TopHat-Cufflinks pipeline provided by the RNASeq app v2.2.10 of the GPRO suite (Hafez *et al*, 2023).

#### Network analyses

Differentially expressed genes related to resulting enriched metabolic pathways (265) were used to assess the enrichment of protein-protein interactions (PPI) and other functional associations. For this purpose, we used the STRING V12 web server (Szklarczyk *et al*, 2023) and seven active interaction sources (experiments, databases, co-expression, text mining, neighborhood, gene fusion, and co-occurrence) and a minimum required interaction score of 0.900.

### Gene over-expression or silencing in GIST

MITF was overexpressed using MITF A GFPpcDNA 3.1+ /C-EGFP, and pEGFP N3 was used as the control plasmid (Genscript, Oxford, UK). pEGFP-LC3 and dual-fluorescene mRFP-GFP-LC3 plasmids were gifts from Tamotsu Yoshimori (Addgene plasmid # 21073 and # 21074) (Kabeya, 2000; Kimura *et al*, 2007).GFP plasmids were transfected into GIST-T1 cells using Lipofectamine LTX (Invitrogen) following the manufacturer’s instructions with slight modifications. Relation (2,5 µg) DNA +13,75 µL lipofectamine/ 2mL). Lentiviral particles to silence the MITF gene expression were previously described elsewhere(Proaño-Pérez *et al*, 2023b). Infected cells were cultured with puromycin (1µg/ml). NT shRNA or MITF shRNA GIST-T1 cells were transiently transfected with pEGFP-LC3. Dual-fluorescence mRFP-GFP-LC3 stable transfectants were produced using Lipofectamine LTX, as described above and selected with G481 (400µg/ml) for 4 weeks. Dual-fluorescence mRFP-GFP-LC3GFP stable transfectants were infected with NT shRNA or MITF shRNAs and maintained with puromycin (1µg/ml) and G418 (400 µg/ml).

### Western blotting

Cellular fractioning was performed as described elsewhere (Suzuki *et al*, 2010). Western blotting was conducted as described (Serrano-Candelas *et al*, 2018). According to the manufacturer’s recommendations, the total protein concentrations were determined using the ProteinAssay Dye Bio-Rad Kit (Bio-Rad Laboratories (RRID:SCR_008426), Inc. Hercules, CA, USA). Electrophoresis and protein blotting was performed using NuPageTM 4–12% and 12% Bis-Tris Gel 1.5 mm * 15 w (Invitrogen, Waltham, MA, USA); after that, electrotransferred to polyvinylidene difluoride (PVDF) membranes (Millipore, Bedford, MA, USA). Membranes were blocked and incubated with primary antibodies. After TTBS washes, membranes were incubated with secondary anti-mouse or anti-rabbit antibodies. Proteins were visualized by enhanced chemiluminescence (WesternBrightTM ECL, Advansta, San Jose, CA, USA).

### Autophagy experiments

Cells were starved overnight with buffer EBSS (B32750, R& D Systems, Minneapolis, MN USA) or treated for 16h with mTOR inhibitor torin1 (200 nM)(Sigma Aldrich, St Louis, MO, USA) and protease inhibitors, pepstatin A (10 ug/mL), and E64D (10 µg/mL) (Sigma-Aldrich). Autophagy induction was performed in GIST-T1 and after overexpressing or silencing MITF.

#### MITF overexpression

MITF-A-GFP or GFP transiently transfected for 24 hours were treated with torin 1 (or DMSO-vehicle) and protease inhibitors, pepstatin A, and E64D for 16 hours. MITF-GFP and LC3II expression was analyzed by western blot.

#### MITF silencing

GIST-T1 were transduced with NT shRNA, MITF shRNA-1, or MITF shRNA-3 lentiviral particles and maintained with puromycin 1ug/mL to select transduced cells. On the 5th day post-transduction, cells were transfected with pEGFP-LC3 and divided for protein expression analysis in a 6-well plate (1×10^6^ cells/w) and resuspended (0.1×10^6^ cells/w) in a 24-well plate on cover slides. After that, at six days post-transduction, cells in both well-plates were treated with torin1 (200nM) +/- pepstatin A (10ug/mL) and E64D (10ug/mL) for 16h. Cells that grew on coverslips were fixed, and GFP-LC3 expression was checked under the microscope. Cells that grew in the p6-plate were lysed for western blot assays.

An RFP-GFP-LC3 GIST-T1 stable cell line was obtained to analyze autophagy flux. The expression of GFP and RFP was monitored for four weeks. Cells were transduced with NT shRNA or MITF shRNA-1, -3 and maintained with puromycin (1µg/ml) and G418 (400 µg/ml). On the 5th day, cells were harvested and divided for protein expression analysis in a conventional 6-well plate (1×10^6^ cells/w) and resuspended (0.1×10^6^ cells/w) in a 24-well plate on cover slides for immunofluorescence studies at six days cells were treated with torin 1 (200nM) +/- pepstatin A 10µg/mL, and E64D 10µg/mL for 16h. Cells that grew on coverslips were fixed, and cells that grew in a 6-well plate were recollected in a lysis buffer to analyze protein concentration and perform a Western blot.

#### Quantitative GFP-LC3 and RFP-GFP-LC3 analyses

GFP or dual RFP-GFP expression was measured by fluorescence microscopy. For that, cells were fixed with 4% paraformaldehyde (Merck KGaA, Darmstadt, Germany), and nuclei were stained with Hoescht 3342 (Merck KGaA) and finally mounted on a slide with the mountant liquid ProLong Gold Antifade Mountant (Thermo Fisher Scientific). Samples were observed with epi-fluorescent microscopy (Leica AF600) with 60x magnification.

Dots were quantified in GFP-LC3-transfected or RFP-GFP-LC3 stable GIST-T1 cells. A minimum of five dots per cell in GFP-LC3 or RFP-GFP-LC3 stable GIST-T1 were needed to be considered positive cells. Regarding GFP-LC3 transfected cells, at least 150 GFP-positive cells per condition were analyzed, and results were expressed as a percentage of GFP-LC3 dot-positive cells/the total number of transfected cells. For RFP-GFP-LC3 tests, at least 30 RFP-GFP-positive GIST-T1 cells for each experimental condition were analyzed. Autophagosomes showed both RFP and GFP signals, while autolysosomes were defined as RFP-positive dots (Ozturk *et al*, 2019). The number of autolysosomes was calculated as follows: Autolysosomes= RFP/positive-dot – GFP/positive dots.

### Isolation of extracellular vesicles from culture media

#### Extracellular vesicle isolation and characterization

Extracellular vesicles (EVs) were isolated from the culture media of lentiviral transduced NT shRNA or MITF shRNA-1 or -3 GIST-T1 by size-exclusion chromatography (SEC) as previously described protocol (Monguió-Tortajada *et al*, 2019a). Briefly, cells were cultured in DMDM supplemented with 10% FBS. On the 4th day after lentiviral transduction, this media was replaced with DMDM and supplemented with EV-depleted FBS. EV-depleted FBS was obtained through ultracentrifugation at 4°C, 100.000 g for 18 h. After 72 h of culture, conditioned media (CM) was collected and subsequently centrifugated at 500 g 4°C for 5 min and at 2000g 4°C for 15 min. This CM was further concentrated using Amicon Ultra-15 100-kDa ultrafilter (UF) units (Millipore, #UFC910024), obtaining 2ml of concentrated conditioned media (CCM) that was stored until further use. In parallel, the remaining cells were counted, and viability was checked using trypan blue staining. Cells were treated for protein extraction (Lysis buffer and stored at -80°C).

EVs were isolated from the CCM using size-exclusion chromatography (SEC), as reported elsewhere(Gámez-Valero *et al*, 2016; Monguió-Tortajada *et al*, 2019a). Sepharose-CL2B (Sigma Aldrich, St Louis, MO, USA) was stacked in a puriflash column Dry Load Empty 12g (20/pk) from Interchim (France)-Cromlab, S.L. (Barcelona, Spain). Five hundred µl/tube fractions were recollected (34 tubes 1.5mL) using PBS1x as elution buffer. Fractions obtained after SEC were analyzed for protein content by measuring their absorbance at 280 nm in a Thermo Scientific Nanodrop® ND-100 (Thermo Fisher Scientific, Waltham, MA).

The SEC fractions showing a low/minimal protein concentration were selected for the forthcoming EV analyses as described (Gámez-Valero *et al*, 2016; Monguió-Tortajada *et al*, 2019b). The presence of specific EV-markers CD9, CD81, and CD63 was determined in these fractions by flow cytometry (FACSDiva (RRID:SCR_001456) Version 6.1.3, BD Biosciences, New Jersey, USA). Mean fluorescence intensity (MFI) values were plotted (Flow Jo software (RRID:SCR_008520), Tree Star, Ashland, OR), and tetraspanin-positive fractions with the highest MFI (fractions 7–12 from our SEC column) were considered EVs enriched fractions and pooled for the subsequent analysis.

EV-enriched pools were characterized by nanoparticle tracking analysis (NTA) and cryo-electron microscopy (cryo-EM), which obtained a rough concentration and checked for morphology and size. Samples were analyzed at Services Technical Scientific of Universitat Autonoma de Barcelona.

The size distribution of EVs was further quantified by measuring the diameter of vesicles in each condition using ImageJ (RRID:SCR_003070).

#### EVs Protein Extraction and Western Blot Analysis

Western blot was used to assess EV and non-EV markers in the EV pools. For this purpose, the protein concentration of EVs was measured using the Micro BCA Protein Assay Kit (Thermo Fisher Scientific). An equal amount of each sample (20ug/25uL) was mixed with reducing 5X Pierce Lane Marker Reducing Sample Buffer (Thermo Fisher Scientific, boiled for 5 min at 95°C, and subjected to western blot as mentioned above.

#### Label-free quantitative proteomics

Purified exosomes were resuspended in Lysis Buffer containing 7M urea, 2 M thiourea, and 50 mM DTT. Protein extracts were diluted in a Laemmli sample buffer and loaded into a 1,5 mm thick polyacrylamide gel with a 4% stacking gel cast over a 12.5% resolving gel. The run was stopped when the front entered 3 mm into the resolving gel so that the whole proteome became concentrated in the stacking/resolving gel interface. Bands were stained with Coomassie Brilliant Blue, excised from the gel, and protein enzymatic cleavage was carried out with trypsin (Promega; 1:20, w/w) at 37 °C for 16 h as previously described (Shevchenko *et al*, 2006). Purification and concentration of peptides were performed using C18 Zip Tip Solid Phase Extraction (Millipore).

#### Liquid chromatography-mass spectrometry (LC-MS/MS)

Peptide mixtures were separated by reverse phase chromatography using an UltiMate 3000 UHLPC System (Thermo Scientific) fitted with an Aurora-packed emitter column (Ionopticks, 25 cm x 75 µm ID, 1.6 µm C18). Samples were first loaded for desalting and concentration into an Acclaim PepMap column (ThermoFisher, 0,5 cm x 300 µm ID, 5 µm C18) packed with the same chemistry as the separating column. Mobile phases were 100% water, 0.1% formic acid (FA) (buffer A), and 100% Acetonitrile 0.1% FA (buffer B). Column gradient was developed in a 120-minute, two-step gradient from 5% B to 20% B in 90 minutes and 20%B to 32% B in 30 minutes. The column was equilibrated in 95% B for 10 minutes and 5% B for 20 minutes. During all processes, the precolumn was in line with the column, and flow was maintained all along the gradient at 300 nl/min. The column temperature was maintained at 40 °C using an integrated column oven (PRSO-V2, Sonation, Biberach, Germany) and interfaced online with the Orbitrap Exploris 480 MS. Spray voltage was set to 2 kV, funnel RF level at 40, and heated capillary temperature at 300 °C. For DDA experiments, full MS resolutions were set to 1200,000 at m/z 200, and the full MS AGC target was set to Standard with an IT mode Auto. The mass range was set to 375–1500. AGC target value for fragment spectra was set to Standard with a resolution of 15,000 and 3 seconds for cycle time. The intensity threshold was kept at 8E3. Isolation width was set at 1.4 m/z. The normalized collision energy was set at 30%. All data were acquired in centroid mode using positive polarity, peptide match was set to off, and isotope exclusion was on.

### Bioinformatic Analyses Proteomics

Raw files were processed with MaxQuant (RRID:SCR_014485) (Cox & Mann, 2008) v1.6.17.0 using the integrated Andromeda Search engine (Cox *et al*, 2011). All data were searched against a target/decoy version of the human Uniprot Reference Proteome (RRID:SCR_002380). The first search peptide tolerance was set to 20 ppm, and the main search peptide tolerance was set to 4.5 ppm. Fragment mass tolerance was set to 20 ppm. Trypsin was specified as the enzyme, cleaving after all lysine and arginine residues and allowing up to two missed cleavages. Carbamidomethylation of cysteine was specified as fixed modification, and peptide N-terminal acetylation, oxidation of methionine, deamidation of asparagine, and glutamine and pyro-glutamate formation from glutamine and glutamate were considered variable modifications with a total of 2 variable modifications per peptide. “Maximum peptide mass” was set to 7500 Da, the “modified peptide minimum score” and “unmodified peptide minimum score” were set to 25, and everything else was set to the default values, including the false discovery rate limit of 1% on both the peptide and protein levels. The obtained quantitative data were exported to Perseus software (version 1.6.15.0)(Tyanova *et al*, 2016) for statistical analysis and data visualization. An unpaired Student’s t-test was used to compare groups directly for total protein analysis. Statistical significance was set at a p-value lower than 0.05 in all cases, and a 1% FDR threshold was considered.

Differentially expressed proteins were considered significant when their Log2 FoldChange was <=-0.38 (downregulated proteins) or >=0.38(up-regulated proteins).

GSEA for all annotated proteins ranked by their T-test statistic was conducted using gseGO and gseKEGG functions of clusterProfiler (RRID: SCR_016884) R package (v.4.8.3) (Wu *et al*, 2021). Data visualization of the clustered top terms of GSEA was performed using the treeplot function in the enrichplot R package (v. 1.20.3)(Visualization *et al*, 2024)

Afterwards, a more restrictive filter was performed. Proteins represented by at least 25 peptides and q-value <0.05 y logFC +- 0,5849 were filtered and described in a heatmap using the heatmap R package (v.1.0.12)(Kolde, 2022; R: The R Project for Statistical Computing). ORA analysis on up-regulated and downregulated proteins was performed using enrichGO and enrichKEGG of clusterProfiler (v.4.8.3) R package. ORA analysis results were represented by the dot plot function from the enrichplot (v. 1.20.3) R package.

## ACKNOWLEDGMENTS

We are indebted to the service of advanced optical microscopy of the CCIT (University of Barcelona) and to the Cytomics core facility of the Institut d’Investigacions Biomèdiques August Pi i Sunyer (IDIBAPS) for technical support. Biorender was used in some figures.

## AUTHOR CONTRIBUTION

E. P-P, M.G., E. S-C, C.S., and M.M. contributed to the study’s conceptualization, data curation, and formal analysis. E. P-P, A.G-V., M.H-L., and E.M. contributed to the methodology, supervision, and validation. C.LL, B.S., D.G-P., and E. S-C. conducted the interpretation of the ChIP-seq, transcriptomic, and proteomic data processed by bioinformatics. E. P-P and M.M. were responsible for writing the original draft. All authors contributed to writing—review, and editing. M.M. and C.S. secured funding acquisition.

## DATA AVAILABILITY

Raw data have been deposited at the NCBI SRA archive with BioProject RRID: SCR_004801) PRJNA748541 and BioSample records (SAMN20336602, SAMN20336603, SAMN20336604, SAMN20336605).

## REFERENCES

Abels ER & Breakefield XO (2016) Introduction to Extracellular Vesicles: Biogenesis, RNA Cargo Selection, Content, Release, and Uptake. Cell Mol Neurobiol 36: 301–312

Andrews S & others (2019) FastQC: a quality control tool for high throughput sequence data. 2010. Https://WwwBioinformaticsBabrahamAcUk/Projects/Fastqc/: http://www.bioinformatics.babraham.ac.uk/projects/

Atay S, Banskota S, Crow J, Sethi G, Rink L & Godwin AK (2014) Oncogenic KIT-containing exosomes increase gastrointestinal stromal tumor cell invasion. Proc Natl Acad Sci 111: 711–716

Atay S, Wilkey DW, Milhem M, Merchant M & Godwin AK (2018) Insights into the proteome of gastrointestinal stromal tumors-derived exosomes reveals new potential diagnostic biomarkers. Mol Cell Proteomics 17: 495–515

Bahrami A, Bianconi V, Pirro M, Orafai HM & Sahebkar A (2020) The role of TFEB in tumor cell autophagy: Diagnostic and therapeutic opportunities. Life Sci 244: 117341

Bauer S, Yu LK, Demetri GD & Fletcher JA (2006) Heat Shock Protein 90 Inhibition in Imatinib-Resistant Gastrointestinal Stromal Tumor. 9153–9161

Blay JY, Kang YK, Nishida T & von Mehren M (2021) Gastrointestinal stromal tumours. Nat Rev Dis Prim 7

Boichuk S, Parry JA, Makielski KR, Litovchick L, Baron JL, Zewe JP, Wozniak A, Mehalek KR, Korzeniewski N, Seneviratne DS, et al (2013) The DREAM complex mediates GIST cell quiescence and is a novel therapeutic target to enhance imatinib-induced apoptosis. Cancer Res 73: 5120–5129

Casali PG, Abecassis N, Bauer S, Biagini R, Bielack S, Bonvalot S, Boukovinas I, Bovee JVMG, Brodowicz T, Broto JM, et al (2018) Gastrointestinal stromal tumours: ESMO–EURACAN Clinical Practice Guidelines for diagnosis, treatment and follow-up. Ann Oncol 29: iv68–iv78

Cortes CJ & La Spada AR (2019) TFEB dysregulation as a driver of autophagy dysfunction in neurodegenerative disease: Molecular mechanisms, cellular processes, and emerging therapeutic opportunities. Neurobiol Dis 122: 83

Cox J & Mann M (2008) MaxQuant enables high peptide identification rates, individualized p.p.b.-range mass accuracies and proteome-wide protein quantification. Nat Biotechnol 26: 1367–1372

Cox J, Neuhauser N, Michalski A, Scheltema RA, Olsen J V. & Mann M (2011) Andromeda: A peptide search engine integrated into the MaxQuant environment. J Proteome Res 10: 1794–1805

Gámez-Valero A, Monguió-Tortajada M, Carreras-Planella L, Franquesa M, Beyer K & Borràs FE (2016) Size-Exclusion Chromatography-based isolation minimally alters Extracellular Vesicles’ characteristics compared to precipitating agents. 6: 1–9

Gao Z, Li C, Sun H, Bian Y, Cui Z, Wang N, Wang Z, Yang Y, Liu Z, He Z, et al (2023) N6-methyladenosine-modified USP13 induces pro-survival autophagy and imatinib resistance via regulating the stabilization of autophagy-related protein 5 in gastrointestinal stromal tumors. Cell Death Differ 30: 544–559

Goding CR & Arnheiter H (2019) MITF—the first 25 years. Genes Dev 33: 983– 1007

Goff L, Trapnell C & Kelley D (2013) Cummerbund: Analysis, Exploration, Manipulation, and Visualization of Cufflinks High. Throughput Seq Data R Packag Version 2

Gozuacik D & Kimchi A (2004) Autophagy as a cell death and tumor suppressor mechanism. Oncogene 23: 2891–2906

Graner MW, Schnell S & Olin MR (2018) Tumor-derived exosomes, microRNAs, and cancer immune suppression. Semin Immunopathol 40: 505–515

Gupta A, Roy S, Lazar AJFF, Wang W-L, McAuliffe JC, Reynoso D, McMahon J, Taguchi T, Floris G, Debiec-Rychter M, et al (2010) Autophagy inhibition and antimalarials promote cell death in gastrointestinal stromal tumor (GIST). Proc Natl Acad Sci 107: 14333–14338

Hafez AI, Soriano B, Elsayed AA, Futami R, Ceprian R, Ramos-Ruiz R, Martinez G, Roig FJ, Torres-Font MA, Naya-Catala F, et al (2023) Client Applications and Server-Side Docker for Management of RNASeq and/or VariantSeq Workflows and Pipelines of the GPRO Suite. Genes (Basel) 14: 267

Hemesath TJ, Steingrímsson E, McGill G, Hansen MJ, Vaught J, Hodgkinson CA, Arnheiter H, Copeland NG, Jenkins NA & Fisher DE (1994) microphthalmia, a critical factor in melanocyte development, defines a discrete transcription factor family. Genes Dev 8: 2770–2780

Hessvik NP, Øverbye A, Brech A, Torgersen ML, Jakobsen IS, Sandvig K & Llorente A (2016) PIKfyve inhibition increases exosome release and induces secretory autophagy. Cell Mol Life Sci 73: 4717–4737

Irazoqui JE (2020) Key Roles of MiT Transcription Factors in Innate Immunity and Inflammation. Trends Immunol 41: 157–171

Jahangiri B, Saei AK, Obi PO, Asghari N, Lorzadeh S, Hekmatirad S, Rahmati M, Velayatipour F, Asghari MH, Saleem A, et al (2022) Exosomes, autophagy and ER stress pathways in human diseases: Cross-regulation and therapeutic approaches. Biochim Biophys Acta - Mol Basis Dis 1868: 166484

Kabeya Y (2000) LC3, a mammalian homologue of yeast Apg8p, is localized in autophagosome membranes after processing. EMBO J 19: 5720–5728

Kawakami A & Fisher DE (2017) The master role of microphthalmia-associated transcription factor in melanocyte and melanoma biology. Lab Investig 97: 649–656

Kim D, Pertea G, Trapnell C, Pimentel H, Kelley R & Salzberg SL (2013) TopHat2: Accurate alignment of transcriptomes in the presence of insertions, deletions and gene fusions. Genome Biol 14: R36

Kim DK, Cho YE, Komarow HD, Bandara G, Song BJ, Olivera A & Metcalfe DD (2018) Mastocytosis-derived extracellular vesicles exhibit a mast cell signature, transfer KIT to stellate cells, and promote their activation. Proc Natl Acad Sci U S A 115: E10692–E10701

Kimura S, Noda T & Yoshimori T (2007) Dissection of the Autophagosome Maturation Process by a Novel Reporter Protein, Tandem Fluorescent-Tagged LC3. Autophagy 3: 452–460

Klionsky DJ, Abdel-Aziz AK, Abdelfatah S, Abdellatif M, Abdoli A, Abel S, Abeliovich H, Abildgaard MH, Abudu YP, Acevedo-Arozena A, et al (2021) Guidelines for the use and interpretation of assays for monitoring autophagy (4th edition)1. Autophagy 17: 1–382

Kolde R (2022) Package ‘pheatmap’: Pretty heatmaps. R Packag: 1–8

Li H & Durbin R (2010) Fast and accurate long-read alignment with Burrows– Wheeler transform. Bioinformatics 26: 589–595

Li J, Dang Y, Gao J, Li Y, Zou J & Shen L (2015) PI3K/AKT/mTOR pathway is activated after imatinib secondary resistance in gastrointestinal stromal tumors (GISTs). Med Oncol 32: 111

Li X, He S & Ma B (2020) Autophagy and autophagy-related proteins in cancer. Mol Cancer 19: 12

Lin J, Li J, Huang B, Liu J, Chen X, Chen X-M, Xu Y-M, Huang L-F & Wang X-Z (2015) Exosomes: novel biomarkers for clinical diagnosis. ScientificWorldJournal 2015: 657086

Löffek S, Zigrino P & Mauch C (2006) Tumorstroma interactions: their role in the control of tumor cell invasion and metastasis. JDDG J der Dtsch Dermatologischen Gesellschaft 4: 496–502

Ma X-H, Piao S, Wang D, Mcafee QW, Nathanson KL, Lum JJ, Li LZ & Amaravadi RK (2011) Measurements of Tumor Cell Autophagy Predict Invasiveness, Resistance to Chemotherapy, and Survival in Melanoma. Clin Cancer Res 17: 3478–3489

Martin FJ, Amode MR, Aneja A, Austine-Orimoloye O, Azov AG, Barnes I, Becker A, Bennett R, Berry A, Bhai J, et al (2023) Ensembl 2023. Nucleic Acids Res 51: D933–D941

Martin M (2011) Cutadapt removes adapter sequences from high-throughput sequencing reads. EMBnet.journal 17: 10

Martina JA, Diab HI, Lishu L, Jeong-A L, Patange S, Raben N & Puertollano R (2014) The nutrient-responsive transcription factor TFE3 promotes autophagy, lysosomal biogenesis, and clearance of cellular debris. Sci Signal 7: ra9

Mizushima N (2007) Autophagy: process and function. Genes Dev 21: 2861– 2873

Monguió-Tortajada M, Gálvez-Montón C, Bayes-Genis A, Roura S & Borràs FE (2019a) Extracellular vesicle isolation methods: rising impact of size-exclusion chromatography. Cell Mol Life Sci

Monguió-Tortajada M, Morón-Font M, Gámez-Valero A, Carreras-Planella L, Borràs FE & Franquesa M (2019b) Extracellular-Vesicle Isolation from Different Biological Fluids by Size-Exclusion Chromatography. Curr Protoc Stem Cell Biol 49

Ngeow KC, Friedrichsen HJ, Li L, Zeng Z, Andrews S, Volpon L, Brunsdon H, Berridge G, Picaud S, Fischer R, et al (2018) BRAF/MAPK and GSK3 signaling converges to control MITF nuclear export. Proc Natl Acad Sci 115: E8668–E8677

Ozturk DG, Kocak M, Akcay A, Kinoglu K, Kara E, Buyuk Y, Kazan H & Gozuacik D (2019) MITF-MIR211 axis is a novel autophagy amplifier system during cellular stress. Autophagy 15: 375–390

Peinado H, Alečković M, Lavotshkin S, Matei I, Costa-Silva B, Moreno-Bueno G, Hergueta-Redondo M, Williams C, García-Santos G, Ghajar C, et al (2012) Melanoma exosomes educate bone marrow progenitor cells toward a pro-metastatic phenotype through MET. Nat Med 18: 883–891

Pogenberg V, Ballesteros-Álvarez J, Schober R, Sigvaldadóttir I, Obarska-Kosinska A, Milewski M, Schindl R, Ögmundsdóttir MH, Steingrímsson E & Wilmanns M (2020) Mechanism of conditional partner selectivity in MITF/TFE family transcription factors with a conserved coiled coil stammer motif. Nucleic Acids Res 48: 934

Proaño-Pérez E, Ollé L, Guo Y, Aparicio C, Guerrero M, Muñoz-Cano R & Martin M (2023a) MITF Downregulation Induces Death in Human Mast Cell Leukemia Cells and Impairs IgE-Dependent Degranulation. Int J Mol Sci 24: 3515

Proaño-Pérez E, Serrano-Candelas E, García-Valverde A, Rosell J, Gómez-Peregrina D, Navinés-Ferrer A, Guerrero M, Serrano C, Martín M, Elizabeth PP, et al (2023b) The microphthalmia-associated transcription factor is involved in gastrointestinal stromal tumor growth. Cancer Gene Ther 30: 115–117

Puertollano R, Ferguson SM, Brugarolas J & Ballabio A (2018) The complex relationship between TFEB transcription factor phosphorylation and subcellular localization. EMBO J 37

R: The R Project for Statistical Computing

Raben N & Puertollano R (2016) TFEB and TFE3: Linking Lysosomes to Cellular Adaptation to Stress. Annu Rev Cell Dev Biol 32: 255–278

Rainer J, Gatto L & Weichenberger CX (2019) ensembldb: an R package to create and use Ensembl-based annotation resources. Bioinformatics 35: 3151–3153

Raposo G & Stoorvogel W (2013) Extracellular vesicles: Exosomes, microvesicles, and friends. J Cell Biol 200: 373–383

Repana D, Nulsen J, Dressler L, Bortolomeazzi M, Venkata SK, Tourna A, Yakovleva A, Palmieri T & Ciccarelli FD (2019) The Network of Cancer Genes (NCG): a comprehensive catalogue of known and candidate cancer genes from cancer sequencing screens. Genome Biol 20: 1

Ron D & Walter P (2007) Signal integration in the endoplasmic reticulum unfolded protein response. Nat Rev Mol Cell Biol 8: 519–529

Runwal G, Stamatakou E, Siddiqi FH, Puri C, Zhu Y & Rubinsztein DC (2019) LC3-positive structures are prominent in autophagy-deficient cells. Sci Rep 9: 10147

Rutkowski DT, Arnold SM, Miller CN, Wu J, Li J, Gunnison KM, Mori K, Sadighi Akha AA, Raden D & Kaufman RJ (2006) Adaptation to ER Stress Is Mediated by Differential Stabilities of Pro-Survival and Pro-Apoptotic mRNAs and Proteins. PLoS Biol 4: e374

Salimi L, Akbari A, Jabbari N, Mojarad B, Vahhabi A, Szafert S, Kalashani SA, Soraya H, Nawaz M & Rezaie J (2020) Synergies in exosomes and autophagy pathways for cellular homeostasis and metastasis of tumor cells. Cell Biosci 10: 64

Schmieder R & Edwards R (2011) Quality control and preprocessing of metagenomic datasets. Bioinformatics 27: 863–864

Serrano-Candelas E, Ainsua-Enrich E, Navinés-Ferrer A, Rodrigues P, García-Valverde A, Bazzocco S, Macaya I, Arribas J, Serrano C, Sayós J, et al (2018) Silencing of adaptor protein SH3BP2 reduces KIT/PDGFRA receptors expression and impairs gastrointestinal stromal tumors growth. Mol Oncol 12: 1383–1397

Serrano C, Mariño-Enríquez A, Tao DL, Ketzer J, Eilers G, Zhu M, Yu C, Mannan AM, Rubin BP, Demetri GD, et al (2019) Complementary activity of tyrosine kinase inhibitors against secondary kit mutations in imatinib-resistant gastrointestinal stromal tumours. Br J Cancer 120: 612–620

Settembre C, De Cegli R, Mansueto G, Saha PK, Vetrini F, Visvikis O, Huynh T, Carissimo A, Palmer D, Klisch TJ, et al (2013) TFEB controls cellular lipid metabolism through a starvation-induced autoregulatory loop. Nat Cell Biol 15: 647–658

Shevchenko A, Tomas H, Havli J, Olsen J V. & Mann M (2006) In-gel digestion for mass spectrometric characterization of proteins and proteomes. Nat Protoc 1: 2856–2860

Singh SS, Vats S, Chia AY-Q, Tan TZ, Deng S, Ong MS, Arfuso F, Yap CT, Goh BC, Sethi G, et al (2018) Dual role of autophagy in hallmarks of cancer. Oncogene 37: 1142–1158

La Spina M, Contreras PS, Rissone A, Meena NK, Jeong E & Martina JA (2021) MiT/TFE Family of Transcription Factors: An Evolutionary Perspective. Front Cell Dev Biol 8: 1–22

Steingrímsson E, Copeland NG & Jenkins NA (2004) Melanocytes and the Microphthalmia Transcription Factor Network. Annu Rev Genet 38: 365– 411

Strub T, Giuliano S, Ye T, Bonet C, Keime C, Kobi D, Le Gras S, Cormont M, Ballotti R, Bertolotto C, et al (2011) Essential role of microphthalmia transcription factor for DNA replication, mitosis and genomic stability in melanoma. Oncogene 30: 2319–2332

Sui S, Ma F, Mi L, Gao L, Yu W, Li M, Feng Z, Huang Y & Wang Q (2022) Circ-CCS enhances autophagy during imatinib resistance of gastrointestinal stromal tumor by regulating miR-197-3p/ATG10 signaling. J Cancer Res Ther 18: 1338–1345

Suzuki K, Bose P, Leong-Quong RY, Fujita DJ & Riabowol K (2010) REAP: A two minute cell fractionation method. BMC Res Notes 3: 294

Szklarczyk D, Kirsch R, Koutrouli M, Nastou K, Mehryary F, Hachilif R, Gable AL, Fang T, Doncheva NT, Pyysalo S, et al (2023) The STRING database in 2023: protein-protein association networks and functional enrichment analyses for any sequenced genome of interest. Nucleic Acids Res 51: D638–D646

Theodosakis N, Pagan AD & Fisher DE (2022) The role of MiT/TFE family members in autophagy regulation. 151–159

Théry C, Zitvogel L & Amigorena S (2002) Exosomes: composition, biogenesis and function. Nat Rev Immunol 2: 569–579 doi:10.1038/nri855 [PREPRINT]

Trapnell C, Roberts A, Goff L, Pertea G, Kim D, Kelley DR, Pimentel H, Salzberg SL, Rinn JL & Pachter L (2012) Differential gene and transcript expression analysis of RNA-seq experiments with TopHat and Cufflinks. Nat Protoc 7: 562

Tyanova S, Temu T, Sinitcyn P, Carlson A, Hein MY, Geiger T, Mann M & Cox J (2016) The Perseus computational platform for comprehensive analysis of (prote)omics data. Nat Methods 13: 731–740 doi:10.1038/nmeth.3901 [PREPRINT]

Visualization T, Enrichment F, Version R, The D, Yulab-smu R, Artistic- L, Annotation V, Utf-VE & Bioconductor R (2024) Package ‘ enrichplot ’

Wang S, Chen Y, Li X, Zhang W, Liu Z, Wu M, Pan Q & Liu H (2020) Emerging role of transcription factor EB in mitochondrial quality control. Biomed Pharmacother 128: 110272

Welsh JA, Goberdhan DCI, O’Driscoll L, Buzas EI, Blenkiron C, Bussolati B, Cai H, Di Vizio D, Driedonks TAP, Erdbrügger U, et al (2024) Minimal information for studies of extracellular vesicles (MISEV2023): From basic to advanced approaches. J Extracell Vesicles 13: e12404

Wu T, Hu E, Xu S, Chen M, Guo P, Dai Z, Feng T, Zhou L, Tang W, Zhan L, et al (2021) clusterProfiler 4.0: A universal enrichment tool for interpreting omics data. Innov (Cambridge 2: 100141

Xing H, Tan J, Miao Y, Lv Y & Zhang Q (2021) Crosstalk between exosomes and autophagy: A review of molecular mechanisms and therapies. J Cell Mol Med 25: 2297–2308

Xu K, He Z, Chen M, Wang N, Zhang D, Yang L, Xu Z & Xu H (2020) HIF-1α regulates cellular metabolism, and Imatinib resistance by targeting phosphogluconate dehydrogenase in gastrointestinal stromal tumors. Cell Death Dis 11: 586

Young MD, Wakefield MJ, Smyth GK & Oshlack A (2010) Gene ontology analysis for RNA-seq: accounting for selection bias. Genome Biol 11: R14

Zhang Y, Liu T, Meyer CA, Eeckhoute J, Johnson DS, Bernstein BE, Nussbaum C, Myers RM, Brown M, Li W, et al (2008) Model-based analysis of ChIP-Seq (MACS). Genome Biol 9: R137

